# Hijacking a dedicated entorhinal-hippocampal extinction circuit to remove traumatic memory

**DOI:** 10.1101/2024.05.13.593830

**Authors:** Ze-Jie Lin, Xue Gu, Wan-Kun Gong, Mo Wang, Yan-Jiao Wu, Qi Wang, Xin-Rong Wu, Xin-Yu Zhao, Michael X. Zhu, Lu-Yang Wang, Quanying Liu, Ti-Fei Yuan, Wei-Guang Li, Tian-Le Xu

## Abstract

Effective psychotherapy of post-traumatic stress disorder (PTSD) remains challenging due to the fragile nature of fear extinction, for which ventral hippocampal CA1 (vCA1) region is considered as a central hub. However, neither the core pathway nor the cellular mechanisms involved in implementing extinction are known. Here, we unveil a direct pathway, where layer 2a fan cells in the lateral entorhinal cortex (LEC) target parvalbumin-expressing interneurons (PV-INs) in the vCA1 region to propel low gamma-band synchronization of the LEC-vCA1 activity during extinction learning. Bidirectional manipulations of either hippocampal PV-INs or LEC fan cells suffice fear extinction. Gamma entrainment of vCA1 by deep brain stimulation (DBS) or noninvasive transcranial alternating current stimulation (tACS) of LEC persistently enhances the PV-IN activity in vCA1, thereby promoting fear extinction. These results demonstrate that the LEC-vCA1 pathway forms a top-down motif to empower low gamma-band oscillations that facilitate fear extinction. Finally, application of low gamma DBS and tACS to a mouse model with persistent PTSD shows potent efficacy, suggesting that the dedicated LEC-vCA1 pathway can be hijacked for therapy to remove traumatic memory trace.

**Graphical Abstract:** 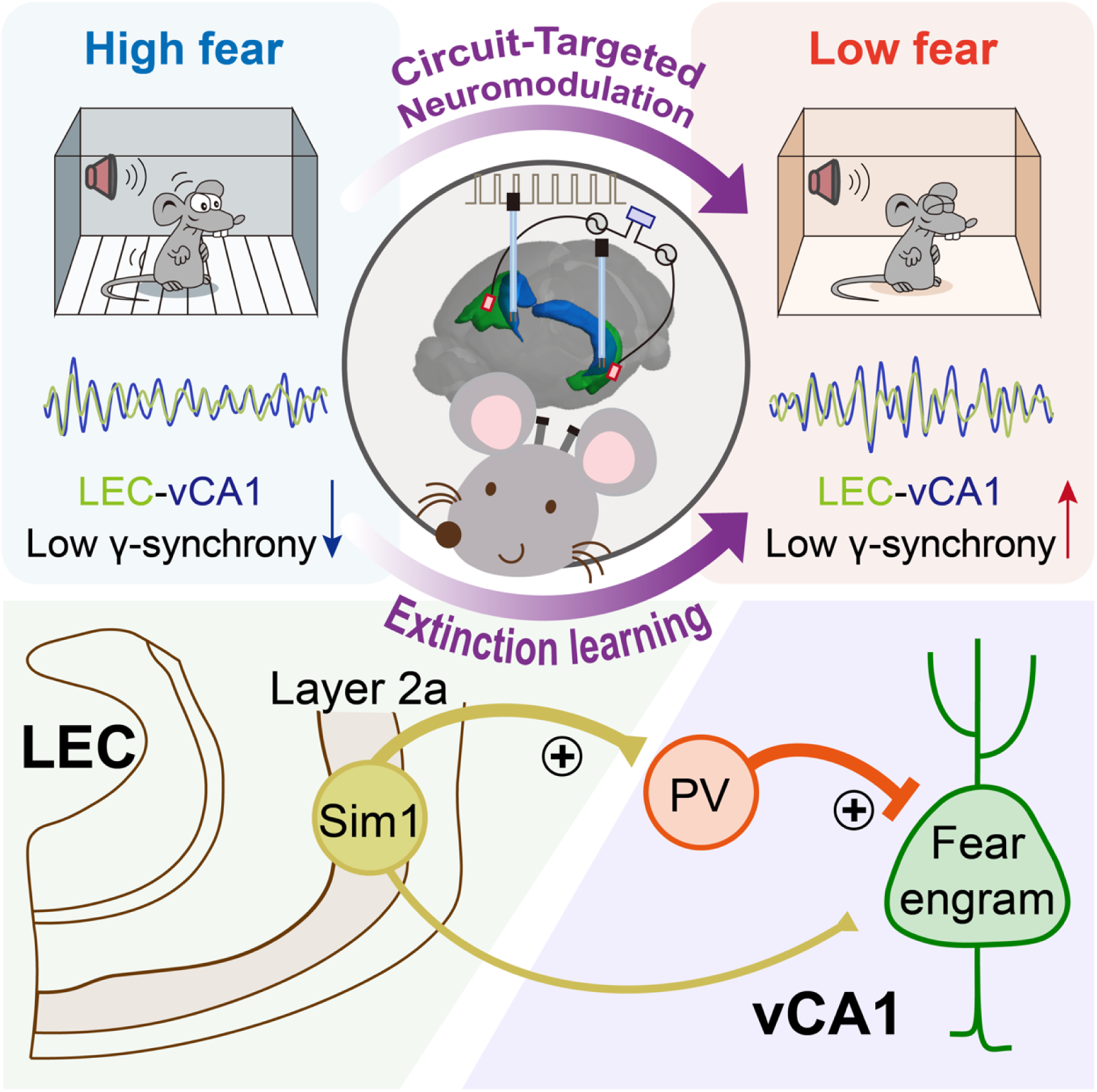

## Introduction

Fear extinction plays a pivotal role in mitigating traumatic memory, facilitating adaptive responses to dynamic environments, and is crucial in psychotherapy for anxiety disorders and post-traumatic stress disorder (PTSD) (1, 2). However, current therapeutic approaches, including drugs and electromagnetic brain stimulations, often lack precision in targets and reliability in outcomes (2). This ambiguity may stem from a limited mechanistic understanding of fear extinction, hindering the development of circuit- and cell-specific interventions. Fear extinction primarily relies on tripartite cortical-subcortical neural circuits, including medial prefrontal cortex (mPFC), basolateral amygdala (BLA), and hippocampus (3–7). Yet, the core pathway and cellular mechanisms governing this tripartite circuitry that drives fear extinction remain elusive. Identifying and harnessing key top-down circuit motifs and cellular ensembles inherent in the natural extinction process holds promise for developing neuromodulation strategies to target pathway- and cell-specific circuits for PTSD treatment.

The hippocampus (HPC), crucial for declarative memory, receives diverse inputs from the neocortex through parahippocampal structures (8, 9), notably the entorhinal cortex (EC) (10). Structurally and functionally, the HPC is divided into the dorsal (dHPC) and ventral (vHPC) regions, associated with spatial memory and emotional processing, respectively (11). The EC comprises the lateral entorhinal cortex (LEC) and the medial entorhinal cortex (MEC), linked to object recognition and spatial learning, respectively (12–14). As a major memory hub, the entorhinal-hippocampal system coordinates projections and synchronizes neural oscillations between brain regions. Despite the well- studied entorhinal-dorsal hippocampal network supporting spatial navigation and associative memory (12–19), the connectivity, activity, and behavioral implications of the ventral hippocampal-entorhinal network remain enigmatic.

Circuit oscillations, arising from synchronized or cooperative activities among different neuronal populations, enable fast transitions between large-scale network states (20, 21). The interplay between circuit oscillations, long-term synaptic plasticity, and recruitment of memory engrams shapes the encoding and retrieval of memories (22, 23). Retrieval of fear memory correlates with amygdalar and hippocampal theta rhythm synchronization (24). Additionally, expression of fear memory involves oscillatory activity in the 3–6 Hz range within the BLA, along with coherence shifting toward the 3–6 Hz range between the BLA and mPFC (25, 26). Conversely, fear extinction remodels the network of inhibitory interneurons in the BLA, allowing a competition between a 6–12 Hz oscillation and the fear- associated 3–6 Hz oscillation (25, 27). This underscores the significance of local and interregional experience-dependent resonance in governing dynamic expression of fear memory. In parallel, gamma oscillations in the hippocampus enhance sensory processing, attention, and memory (28–31). Pathway- specific gamma oscillations facilitate task-relevant information routing between distinct neuronal subpopulations within the entorhinal-hippocampal circuit (14). These findings suggest that oscillatory activity within the entorhinal-hippocampal circuit may be related to fear extinction, representing a form of inhibitory learning. The circuit organization, along with oscillatory dynamics concerning cell- type specific connectivity between EC and ventral hippocampus involved in fear extinction and its potential for therapeutic neuromodulation of PTSD, remain unexplored.

Our study reveals a novel direct projection from LEC layer 2a fan cells to ventral hippocampal CA1 (vCA1) parvalbumin-expressing interneurons (PV-INs), distinct from established indirect trisynaptic pathways observed from LEC layer 2a to the dorsal hippocampus (13, 19, 32, 33). Further exploration of neural oscillations within the EC-vCA1 network reveals that extinction training is associated with heightened low-gamma rhythms and synchronization between LEC and vCA1 regions. This oscillation is mediated by vCA1 PV-INs directly innervated by LEC layer 2a fan cells. Importantly, entraining the newly identified LEC-vCA1 pathway with clinically available interventions, such as deep brain stimulation (DBS) and transcranial alternating current stimulation (tACS) (34–38), results in a robust attenuation of fear memory. This provides a proof of principle for alleviating traumatic memories using readily available strategies.

## Results

### Fear extinction induces low-gamma rhythm synchronization between LEC and vCA1

To explore the entorhinal cortex-vCA1 functional connectivity in fear extinction, we implanted electrodes in the vCA1, LEC, and MEC to record local field potentials (LFPs) during fear extinction. Mice underwent auditory fear conditioning followed by extinction training, exhibiting a gradual reduction in freezing responses (Figure 1, A–D). LFP analysis revealed increased low-theta (3–6 Hz) but not high-theta (6– 12 Hz) oscillations during early extinction (Early-Ext., CS1-4), paralleling previous findings in the basolateral amygdala (BLA) (25, 26), with vCA1 showing decreased low-gamma power initially and increased power during late extinction (Late-Ext., CS17-20) (Figure 1, E–G). Similar oscillatory changes were observed in LEC and MEC (Supplemental Figure 1), with negative correlations between conditioned freezing and low-gamma power in all regions (Figure 1H and Supplemental Figure 1, D and G). Phase synchronization analysis using the weighted phase lag index (wPLI) showed higher synchrony between LEC-vCA1 low-gamma oscillations during late extinction, highlighting their role in fear extinction compared to MEC-vCA1 synchronization (Figure 1, I–L).

**Figure 1.**
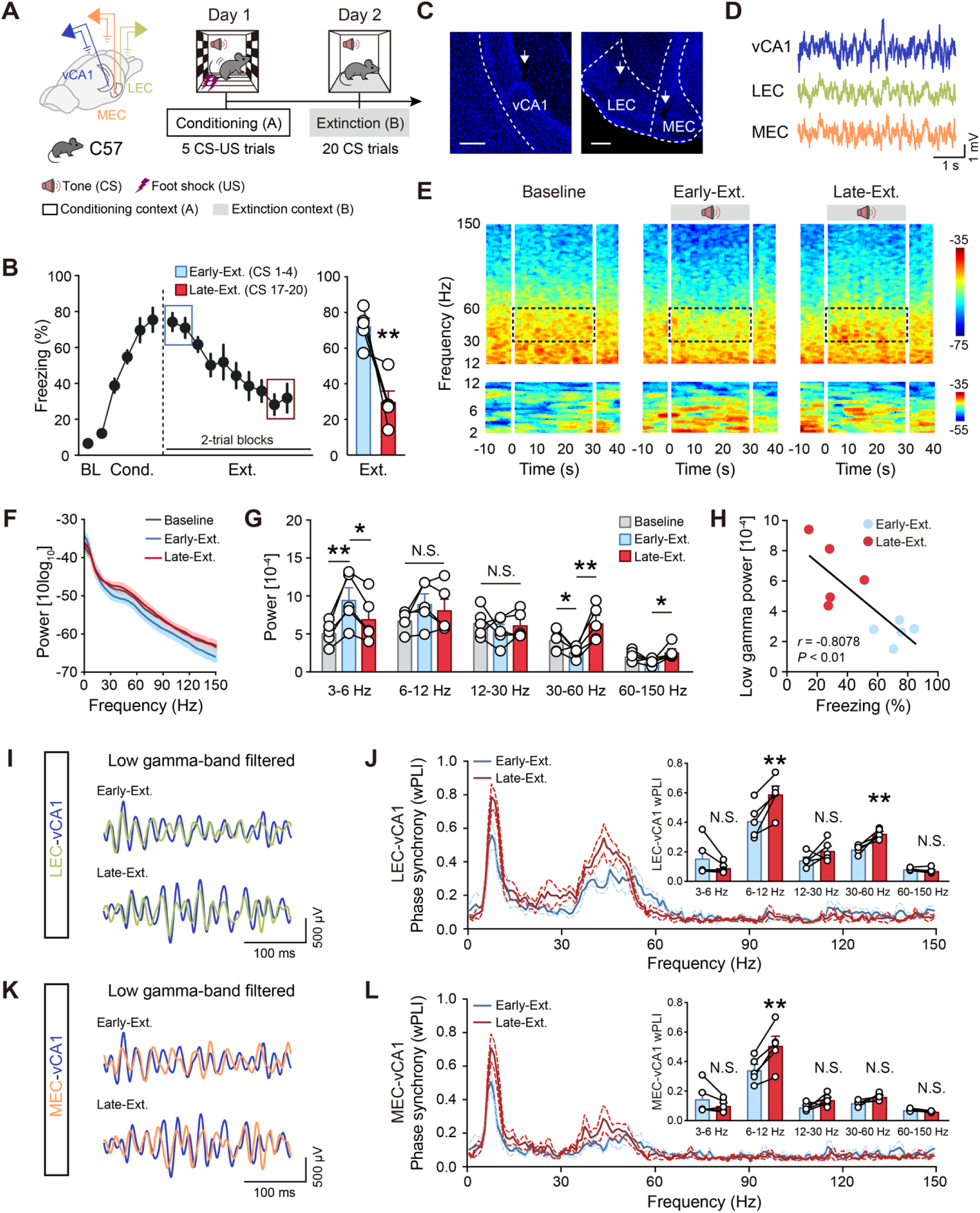
Fear extinction recruits low-gamma oscillatory synchrony between the LEC and vCA1. (**A**) Schematics of electrode implantation and experimental design for mice subject to fear conditioning (day 1, context A) and extinction training (day 2, context B). (**B**) Time courses of freezing responses to the CS during fear conditioning and extinction training sessions (left). While the freezing response during conditioning was calculated by percent freezing time to CS in individual trials, those during extinction training were calculated by the average freezing responses of two consecutive trials. The same convention was used for the calculation of all behavioral results. Freezing responses to the CS during early extinction training (CS1-4, referred to as Early-Ext.) and late extinction training (CS17-20, referred to as Late-Ext.) (right). Data are mean ± S.E.M. \*\**P* < 0.01, paired Student’s *t*-test. *n* = 5 mice. (**C**) Representative images showing vCA1, LEC and MEC electrode placements. Scale bar, 200 μm. (**D**) Representative traces of vCA1, LEC and MEC local field potential (LFP) recordings. (**E**) Representative spectrograms of LFP recorded in vCA1 during Baseline (left), Early-Ext. (middle) and Late-Ext.(right) sessions. 0-30 s represents the tone given during the extinction training. (**F**) Power spectrum of vCA1 LFP recordings during Baseline, Early-Ext. and Late Ext.. Solid lines represent the averages and shaded areas indicate SEM. (**G**) Average power of vCA1 LFP recordings during Baseline, Early-Ext. and Late Ext.. Histograms represent mean ± SEM and circles denote individual mice. *n* = 5. N.S., no significant difference, \**P* < 0.05, \*\**P* < 0.01, paired Student’s t-test. (**H**) Linear regression of freezing responses *vs*. vCA1 low-gamma power during Early-Ext. and Late Ext. sessions. (**I**) Examples of low-gamma frequency filtered LEC and vCA1 LFP recordings recorded during Early-Ext. and Late Ext. sessions. (**J**) Extinction training-induced phase synchrony for LEC-vCA1 LFPs weighted phase lag index (wPLI) in the Early-Ext. and Late-Ext. sessions, respectively. The inset shows different phase synchrony quantified using the wPLI between LEC and vCA1 LFPs. Histograms represent mean ± SEM and circles denote individual mice. *n* = 5. N.S., no significant difference, \*\**P* < 0.01, paired Student’s t-test. (**K** and **L**) The same as (**I** and **J**) for MEC-vCA1 LFPs and wPLI. *n* = 5. N.S., no significant difference, \*\**P* < 0.01, paired Student’s t-test.

### Low-gamma synchronization between LEC and vCA1 during fear extinction requires the vCA1

*PV-INs*. Neuronal oscillations result from the dynamic interplay between excitation and inhibition (20, 21, 39), with inhibitory interneurons (40), including PV-INs, somatostatin-expressing interneurons (SST-INs), and vasoactive intestinal peptide-expressing interneurons (VIP-INs), orchestrating synchronized activity in the hippocampus. To identify the interneuron subtype involved in fear extinction-induced network reorganization, we selectively labeled GABAergic neurons in vCA1 using AAV-DIO-mCherry in PV-Cre, SST-Cre, and VIP-Cre mice (Supplemental Figure 2A). Fear extinction selectively activated PV-INs, as indicated by increased PV-mCherry^+^/c-fos^+^ cells compared to SST- INs or VIP-INs (Supplemental Figure 2, B and C). *In vivo* Ca^2+^ recordings using fiber photometry to detect cell-type-specific GCaMP6m fluorescence changes further confirmed the specific activation of PV-INs during fear extinction in vCA1 (Figure 2, A and B and Supplemental Figure 2D). Real-time Ca^2+^ signals revealed increased PV-IN activity during the Late-Ext. phase (Figure 2, C–E), while SST- INs and VIP-INs remained unchanged (Supplemental Figure 2, E–J), reinforcing the specific involvement of PV-INs in fear extinction.

**Figure 2.**
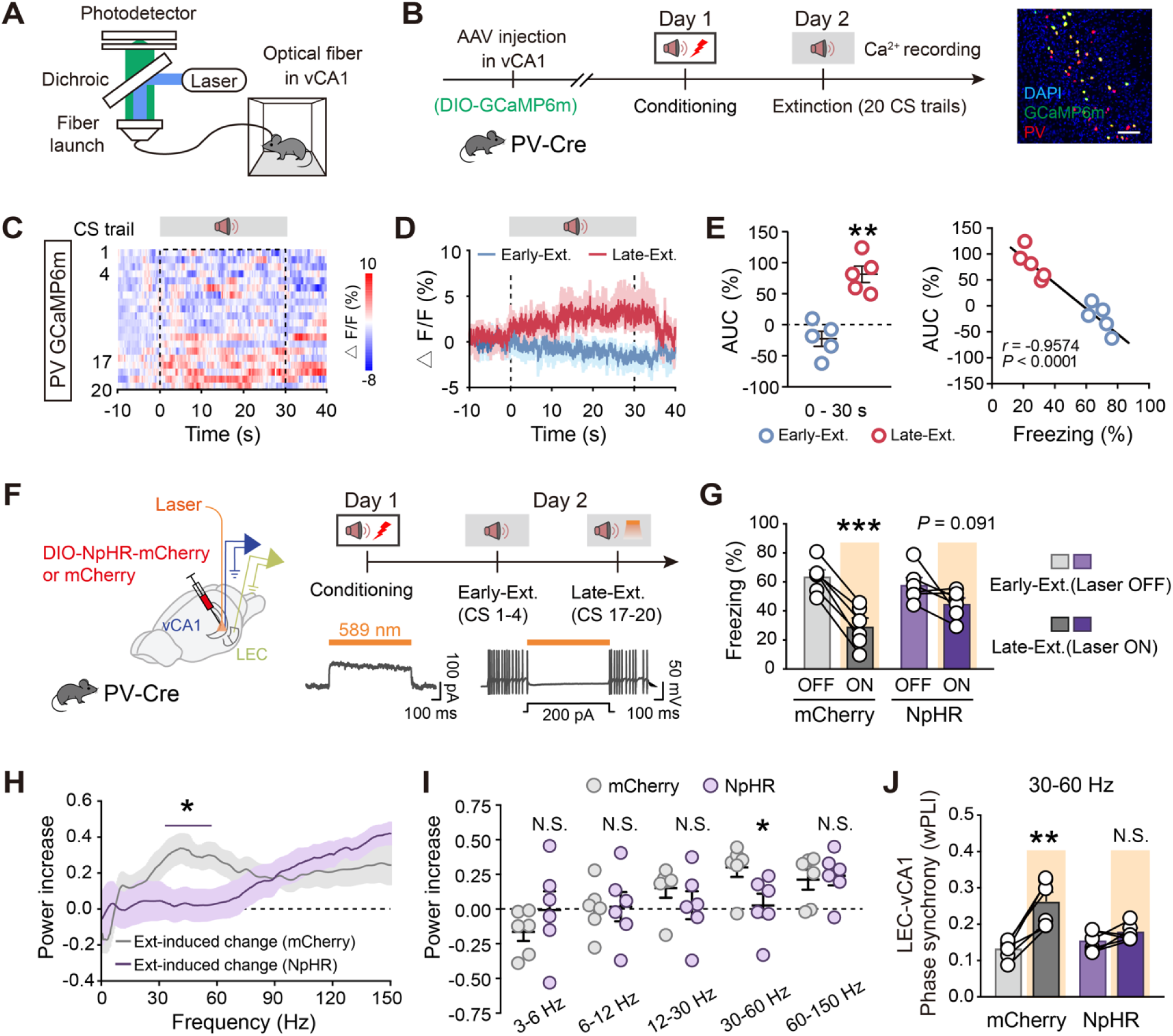
Hyperexcitability of vCA1 PV-INs in the late extinction contributes to LEC-vCA1 low gamma synchronization. (**A**) Schematic illustration of the fiber photometry setup. (**B**) Schematics of AAV injections and experimental design, as well as immunostaining confirming the specificity of GCaMP6m expression in the PV-INs. Scale bar, 100 μm. (**C**) Heatmap of calcium signals in the PV-INs during extinction training sessions. (**D**) Average calcium signals in the PV-INs during Early-Ext. and Late-Ext. Data are mean ± SEM. *n* = 5 mice. (**E**) Activity of the PV-INs (area under the curve, AUC) and correlation of freezing responses with the calcium signals during Early-Ext. and Late-Ext. Quantification of AUC of calcium signals in the PV-INs (left). Data are mean ± SEM. N.S., no significant difference, \**P* < 0.05, \*\**P* < 0.01, paired Student’s t-test. Linear regression of freezing responses *vs*. AUC of PV-INs GCaMP signals during Early-Ext. and Late Ext. (right). (**F**) Schematics of stereotaxic surgery and experimental design. Optogenetic stimulation (yellow light) was delivered during Late-Ext. on Day 2. Traces are representatives of yellow-light-evoked (1 s) outward current and blockade of action potential firing by the yellow light recorded from NpHR-expressing neurons. (**G**) Freezing responses to the CS during Early-Ext. and Late-Ext. mCherry group, *n* = 6 mice; NpHR group, *n* = 6 mice. Data are mean ± SEM. \*\*\**P* < 0.001, paired Student’s *t*-test. (**H**) Extinction-induced changes in power spectrum of vCA1 LFP recordings. Shown are mean ± SEM of power (Late-Ext. - Early-Ext.) / (Late-Ext. + Early-Ext.). *n* = 6 mice per group. Purple line indicates frequencies with a significant effect (\**P* < 0.05 with Bonferroni correction for multiple comparisons). (**I**) Average power increase of vCA1 LFP recordings. Data are mean ± SEM and circles denote individual mice. *n* = 6 mice per group. N.S., no significant difference, \**P* < 0.05, unpaired Student’s t-test. (**J**) Low-gamma phase synchrony quantified using the wPLI between LEC and vCA1 LFPs. Histograms represent mean ± SEM and circles denote individual mice. *n* = 6 mice per group. N.S., no significant difference, \*\**P* < 0.01, paired Student’s t-test.

To evaluate the impact of PV-INs on neural oscillations during fear extinction, we bilaterally injected AAV-DIO-NpHR-mCherry or control virus into the vCA1 of PV-Cre mice and implanted optical fibers targeting vCA1, and placed LFP electrodes in both vCA1 and LEC (Figure 2F). Optical inhibition of vCA1 PV-INs during the Late-Ext. phase resulted in significantly higher freezing compared to the control group (Figure 2G). Additionally, the control group exhibited increased low- gamma oscillations in vCA1 during the Late-Ext. phase (Figure 2H), which were abolished by inhibiting PV-INs, along with disrupted LEC-vCA1 synchronization (Figure 2, I and J). These results emphasize the necessity of vCA1 PV-INs for fear extinction, likely through promoting LEC-vCA1 low-gamma synchronization.

### LEC SIM1^+^ layer 2a fan cells are the main projection neurons to vCA1 PV-INs

To map the monosynaptic inputs to vCA1 PV-INs, we employed Cre-dependent rabies-virus (RV)-mediated retrograde tracing in PV-Cre mice. Starter PV-INs (EGFP^+^ and DsRed^+^) in vCA1 were identified, and DsRed^+^ neurons outside vCA1 served as long-range presynaptic neurons (Figure 3A). PV-INs received inputs from LEC, dorsal hippocampus (dHPC), and medial septal nucleus (MS), with fewer inputs from MEC (Figure 3, B and C and Supplemental Figure 3, B and C). Using AAV with CaMKIIα promoter to express ChR2 in LEC neurons, we recorded light-induced excitatory postsynaptic currents (EPSCs) in vCA1 PV-INs, confirming monosynaptic glutamatergic connections between LEC and vCA1 PV-INs.

**Figure 3.**
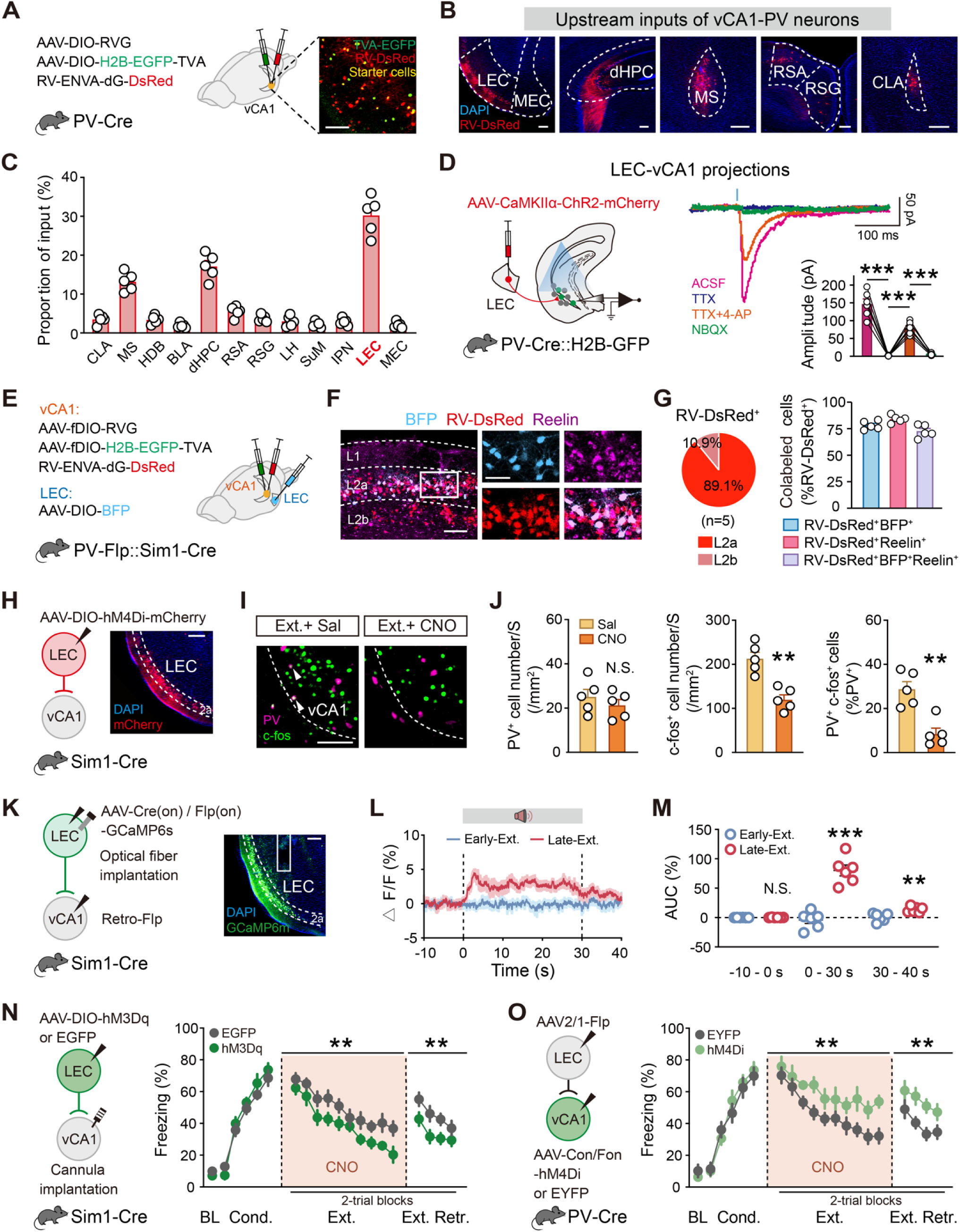
vCA1 PV-INs receive strong excitatory input from Sim1^+^ fan cells in LEC-layer 2a during extinction. (**A**) Schematics of AAV injections and experimental design (left) and a representative image of vCA1 TVA-EGFP and RV-DsRed expression at the injection site (right). Scale bar, 100 μm. (**B**) Representative images of the main upstream inputs of PV-INs. Scale bar, 200 μm. (**C**) Distribution of RV-DsRed-labeled neurons. *n* =5 mice. CLA, claustrum; MS, medial septal nucleus; HDB, nucleus of the horizontal limb of the diagonal band; BLA, basolateral amygdalar nucleus; dHPC, dorsal hippocampus; RSA, retrosplenial agranular cortex; RSG, retrosplenial granular cortex; LH, lateral hypothalamic; SuM, supramammillary nucleus; LPN, the interpeduncular nucleus; LEC, lateral entorhinal cortex; MEC, medial entorhinal cortex. (**D**) Patch clamp recordings of activity of vCA1 PV-INs in brain slices upon optogenetic stimulation of LEC-vCA1 projections (left), showing example traces evoked by blue lights in the presence of ACSF, TTX (1 μM), TTX plus 4-AP (100 μM) and NBQX (10 μM). The blue vertical bar above traces indicates photostimulation. The inset histograms represent mean ± SEM and circles denote individual neurons. \*\*\**P* < 0.001, paired Student’s t-test. *n* = 6 neurons from three mice. (**E**–**G**) LEC layer 2a-vCA1 PV-IN projectors are Sim1^+^ fan cells. (**E**) Schematics of AAV injections. (**F**) Representative images of BFP^+^ (blue), RV-DsRed^+^ (red) and Reelin^+^ (purple) immunofluorescence in LEC. Scale bars, 100 μm (left), 50 μm (right). (**G**) LEC neurons projecting to vCA1 PV-INs are mainly located in layer 2a (left) and are characterized by the expression of Reelin (right). Histograms represent mean ± SEM and circles denote individual mice. *n* = 5. (**H**) Schematics of AAV injections and experimental design (left) and representative image of mCherry expression in LEC-layer 2a (right). CNO was administrated 30 min (i.p.) before extinction training. Scale bar, 200 μm. (**I**) Representative images of PV (purple) and c-fos^+^ (green) immunofluorescence in vCA1. The white arrowheads denote colabeled PV^+^/c-fos^+^ cells. Scale bar, 100 μm. (**J**) The number of activated (PV^+^/c-fos^+^) PV-INs was significantly higher in the Ctrl group than the CNO group. Histograms represent mean ± SEM and circles denote individual mice. Ctrl group, *n* = 5 mice; CNO group, *n* = 5 mice. N.S., no significant difference, \*\**P* < 0.01, unpaired Student’s t-test. (**K**-**M**) Ca^2+^ recording of the LEC-vCA1 pathway during extinction. (**K**) Schematics of AAV injections and fiber implantation (left), with representative images of CGaMP6s expression in LEC- layer 2a (right). Scale bar, 200 μm. (**L**) Average calcium signals during Early-Ext. and Late-Ext. (**M**) Activity of calcium signals (AUC) during Early-Ext. and Late-Ext. Data are mean ± SEM. N.S., no significant difference, \*\**P* < 0.01, \*\*\**P* < 0.001, paired Student’s t-test. *n* = 6 mice. (**N** and **O**) Effects of stimulating LEC-layer 2a → vCA1 projection (**N**) and inhibiting LEC → vCA1 PV-IN projection (**O**) on extinction training and retrieval. Schematics of AAV injections (left). Time courses of freezing responses to the CS during fear conditioning, extinction training and extinction retrieval sessions (right). Statistics are as follows: two-way repeated measures ANOVA, main effect of AAV, (**N**) conditioning, F_1,17_ = 1.157, *P* = 0.2971; extinction training, F_1,17_ = 8.686, *P* = 0.0090; extinction retrieval, F_1,17_ = 9.781, *P* = 0.0061. EGFP group, *n* = 10 mice, hM3Dq group, *n* = 9 mice. (**O**) conditioning, F_1,14_ = 0.1024, *P* = 0.7537; extinction training, F_1,14_ = 14.23, *P* = 0.0021; extinction retrieval, F_1,14_ = 12.46, *P* = 0.0033. EYFP group, *n* = 8 mice, hM4Di group, *n* = 8 mice. Data are mean ± SEM. \*\**P* < 0.01.

The LEC neurons projecting to vCA1 PV-INs were primarily located in the superficial sub-layer 2a (Figure 3B and Supplemental Figure 3B), enriched with reelin-positive fan cells (13, 19, 32, 33). To identify these fan cells, we selectively labeled SIM^+^ layer 2a fan cells receiving retrograde signals from vCA1 PV-INs using an intersectional strategy employing the PV-Flp::Sim1-Cre mice (Figure 3E). Flp-dependent transsynaptically labeled presynaptic neurons (DsRed^+^) were mainly in layer 2a (Figure 3F), with the majority co-expressing DsRed and Cre-dependent blue fluorescent protein (BFP) simultaneously (Figure 3G), confirming that LEC SIM1^+^ layer 2a fan cells are the principal projection neurons to vCA1 PV-INs.

### LEC layer 2a fan cells-vCA1 PV-INs pathway orchestrates their synchronization and fear extinction

To solidify the functional role of LEC layer 2a neurons in activating vCA1 PV-INs during fear extinction, we used chemogenetic inhibition with designer receptors activated only by designer drugs (DREADD) in Sim1-Cre mice. Bilateral injections of AAV-DIO-hM4Di-mCherry into the LEC reduced fear extinction-induced activation of vCA1 PV-INs upon clozapine-N-oxide (CNO) administration compared to the saline control (Figure 3, I and J). Fiber photometry of Ca^2+^ signals in vCA1-projecting LEC SIM1^+^ layer 2a fan cells showed a significant increase in cue-induced activity during the Late-Ext. phase (Figure 3, K–M), contrasting with minimal changes in dHPC-vCA1 or MS- vCA1 pathways (Supplemental Figure 4, A–F).

Furthermore, manipulating LEC SIM1^+^ layer 2a fan cells’ impact on neural oscillations during fear extinction, we bilaterally injected AAV-DIO-NpHR-mCherry into the LEC of Sim1-Cre mice (Supplemental Figure 5, A and B). Silencing LEC layer 2a fan cell input by activating NpHR with light abolished Late-Ext.-associated increases in low-gamma power and synchronization (Supplemental Figure 5, C–E). Additionally, we manipulated LEC SIM1^+^ layer 2a fan cell projection in Sim1-Cre mice by chemogenetics using DREADD hM3Dq (i.e. AAV-DIO-hM3Dq-EGFP, with AAV-DIO-EGFP as a control). Specifically increasing presynaptic activities of the fan cell projection by local perfusion of CNO (1 mM, 200 nl) into the axon projection fields in vCA1 significantly lowered levels of freezing during both extinction training and retrieval compared to the control (Figure 3N). Moreover, by targeting an inhibitory DREADD hM4Di (or a control virus without the hM4Di effector) in a Cre- and Flp-dependent (Cre_on_/Flp_on_) manner into vCA1 PV-INs that receive projections from LEC (with an anterogradely transsynaptic AAV2/1-Flp virus injected into LEC), we chemogenetically inhibited this subpopulation of PV-INs with CNO and observed significant increases in freezing during extinction training and retrieval (Figure 3O). These bidirectional manipulations underscore the necessity and sufficiency of the functional connectivity between LEC SIM1^+^ layer 2a fan cells and vCA1 PV-INs in fear extinction, establishing the LEC-vCA1 pathway as an essential top-down motif.

### DBS selectively recruits PV-INs to entrain vCA1 into low-gamma oscillations to propel fear extinction

Given the direct pathway from LEC to vCA1 governing fear extinction via low gamma entrainment, we explored the efficacy of frequency-dependent deep brain stimulation (DBS) therapy in fear memory mice. We paired CS with DBS at different frequencies (20 Hz, 40 Hz, and 130 Hz, with 40 Hz falling within the low-gamma frequency range) during extinction training. Remarkably, mice exposed to CS-paired 40 Hz DBS exhibited a significant reduction in freezing behavior, persisting into extinction retrieval, compared to those with 20 Hz DBS or without DBS (Figure 4, A–C). Behavioral alterations were specific to fear extinction, as open field and elevated plus maze tests showed no impact on exploratory behaviors or basal anxiety levels (Supplemental Figure 6). Mechanistically, 40 Hz DBS led to much higher activation of PV-INs compared to other frequencies, correlating with behavioral outcomes (Figure 4, D and E). Additionally, chemogenetic inhibition of PV-INs abolished the effects of DBS on fear extinction, while inhibition of other interneuron types did not (Figure 4F and Supplemental Figure 7). Optical stimulation of vCA1 PV-INs mimicked the effects of DBS on fear extinction (Supplemental Figure 8), indicating selective recruitment of PV-INs by 40 Hz DBS for enhanced extinction efficacy.

**Figure 4.**
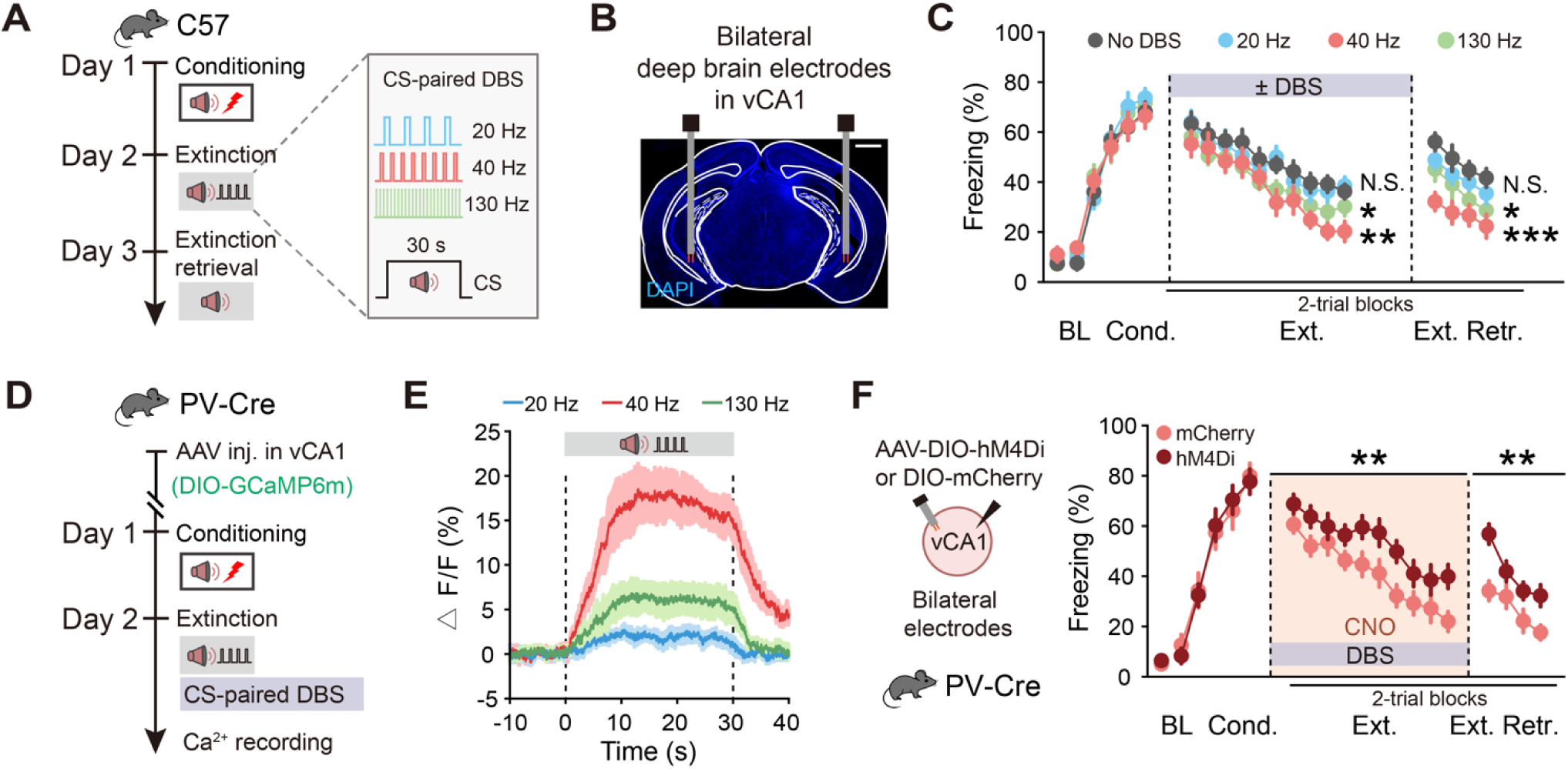
Low-gamma DBS-induced long term extinction promotion depends on the activation of vCA1 PV-INs. (**A**) Schematics of experimental design. CS is paired with DBS of different frequencies (20 Hz, 40 Hz, 130 Hz) during extinction training. (**B**) Representative image showing electrode placements in vCA1. Scale bar, 1 mm. (**C**) Time courses of freezing responses to the CS during fear conditioning, extinction training and extinction retrieval sessions. Statistics are as follows: two-way repeated measures ANOVA, main effect of DBS frequency, conditioning, F_3,34_ = 0.3943, *P* = 0.7579. No DBS *vs.* 20 Hz DBS, extinction training, F_1,16_ = 0.3954, *P* = 0.5383; extinction retrieval, F_1,16_ = 2.126, *P* = 0.1642. No DBS *vs.* 40 Hz DBS, extinction training, F_1,16_ = 12.91, *P* = 0.0024; extinction retrieval, F_1,16_ = 24.91, *P* = 0.0001. No DBS *vs.* 130 Hz DBS, extinction training, F_1,16_ = 5.237, *P* = 0.0360; extinction retrieval, F_1,16_ = 5.192, *P* = 0.0368. No DBS group, *n* = 8 mice, 20 Hz DBS group, *n* = 10 mice, 40 Hz DBS group, *n* = 10 mice, 130 Hz DBS group, *n* = 10 mice. Data are mean ± SEM. N.S., no significant difference, \**P* < 0.05, \*\**P* < 0.01, \*\*\**P* < 0.001. (**D**) Schematics of AAV injections and experimental design. (**E**) Average calcium signals in PV neurons during extinction training paired with DBS of different frequencies. Data are mean ± SEM. 20 Hz group, *n* = 5 mice; 40 Hz group, *n* = 5 mice; 130 Hz group, *n* = 6 mice. (**F**) Effect of inhibiting vCA1 PV-INs on DBS-induced extinction promotion. Time courses of freezing responses to the CS during fear conditioning, extinction training and extinction retrieval sessions. Statistics are as follows: two-way repeated measures ANOVA, main effect of AAV, conditioning, F_1,18_ = 0.0015, *P* = 0.9699; extinction training, F_1,18_ = 12.56, *P* = 0.0023; extinction retrieval, F_1,18_ = 14.80, *P* = 0.0012. mCherry group, *n* = 10 mice, hM4Di group, *n* = 10 mice. Data are mean ± SEM. \*\**P* < 0.01.

### PV-INs with high basal firing rate are preferentially recruited by low-gamma DBS in vCA1

To dissect vCA1 PV-IN firing dynamics during extinction retrieval with high precision, we conducted single-unit electrophysiological recordings. By opto-tagging PV-INs with AAV-DIO-ChR2-mCherry in PV-Cre mice and employing an optrode above the vCA1 injection site (Figure 5A and Supplemental Figure 9A), we captured 503 well-isolated neurons, including 27 optogenetically tagged PV-INs, 409 wide spike neurons (putative pyramidal neurons), and 67 narrow spike neurons (putative interneurons) (Figure 5, B and C and Supplemental Figure 9, B-E). Among the narrow spike population, we identified 49 putative PV-INs including the optogenetically tagged (*n* = 27) and fast-spiking putative (*n* = 22) interneurons, categorized by basal firing rates into high (>30 Hz), medium (15–30 Hz), and low (<15 Hz) groups.

**Fig. 5.**
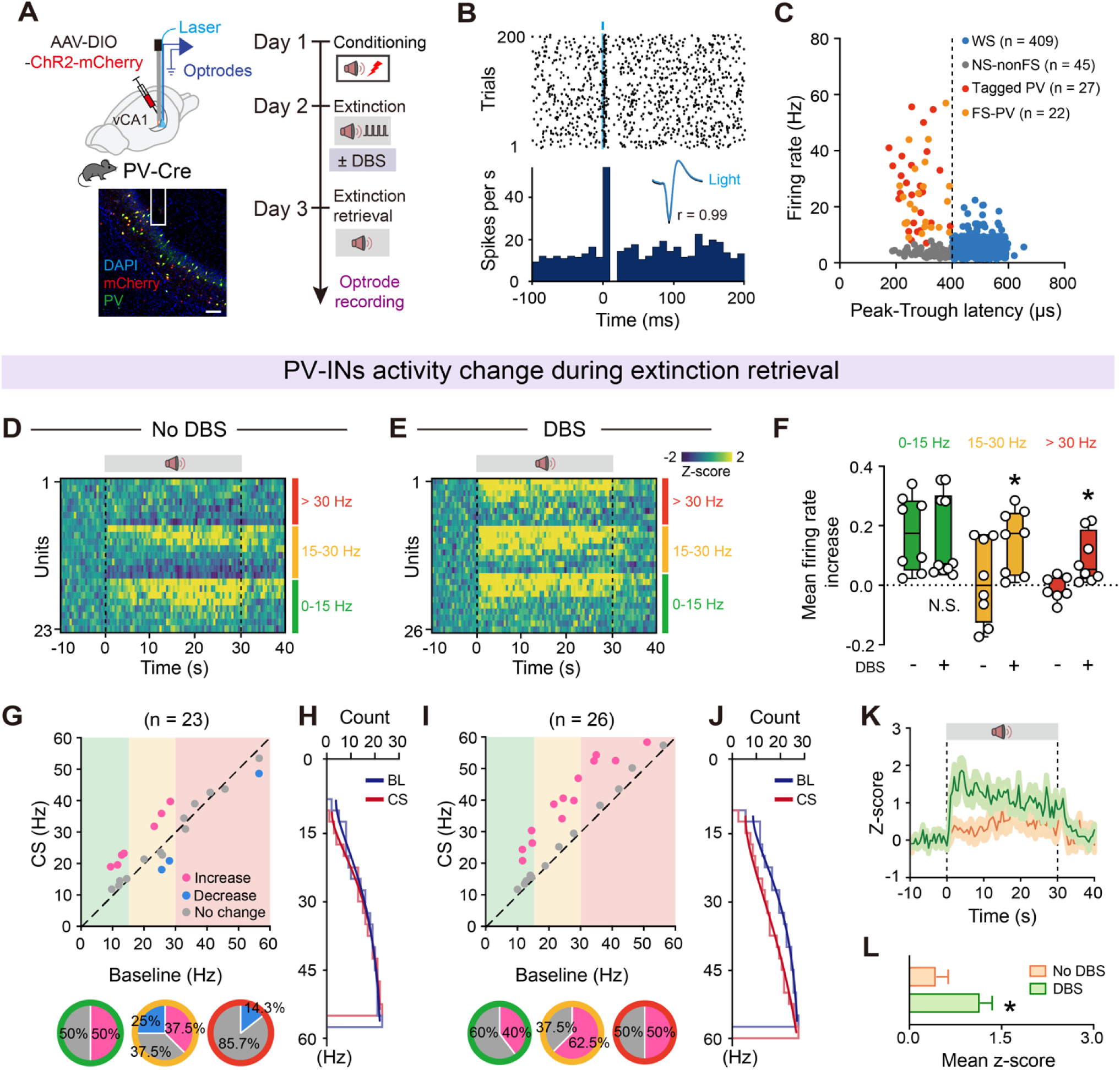
Low-gamma DBS paired extinction training induces sustained activation of high-firing rate vCA1 PV-INs during extinction retrieval. (**A**) Schematics of experimental design (top) and representative image of virus expression (bottom). Scale bar, 100 μm. (**B**) Raster plot (top) and peri-stimulus time histogram (PSTH, bottom) of a representative tagged PV-INs. In the inset, light-evoked spike waveforms (blue) were similar to spontaneous ones (black). Pearson’s correlation, r = 0.99. (**C**) Classification of recorded vCA1 neurons into WS putative pyramidal cells (blue circles), NS-nonFS (gray circles), Tagged PV (red circles) and FS-PV (orange circles) based on peak-to-trough latency and baseline firing rate. (**D** and **E**) Heatmaps showing responses of PV-INs with different baseline firing rate during extinction retrieval. (**D**) No DBS manipulation during extinction training. (**E**) CS was paired with 40 Hz DBS during extinction training. The red parts indicate PV-INs firing at frequencies higher than 30 Hz, the yellow parts indicate firing between 15 and 30 Hz, and the green parts indicate firing between 0 and 15 Hz. (**F**) Box plots of extinction training-induced firing rate changes without and with DBS pairing during extinction retrieval. The center line shows median, box edges indicate top and bottom quartiles, whiskers extend to minimum and maximum values. Circles denote individual neurons. N.S., no significant difference, \**P* < 0.05, unpaired Student’s t-test. (**G** and **H**) Correlation of firing rate at baseline and during CS for individual PV-INs from No DBS manipulation mice. (**G**) Colored circles indicate neurons that showed significant firing rate increase (pink) or decrease (blue) or no significant change (gray). The pie graphs at the bottom show the percentage of neurons (Outer ring color represents PV-INs in different frequency segments) that had significantly higher (pink), lower (blue) or unchanged (gray) firing rates during extinction retrieval. (**H**) Cumulative distributions of firing frequencies for individual PV-INs. Blue line indicates the cumulative plot from the baseline (BL) and red line indicates the cumulative plot obtained during CS. (**I** and **J**) The same as (**G** and **H**) for the correlation of firing rate during BL and CS for individual PV-INs from DBS manipulation mice. (**K** and **L**) Z-scored signal changes of PV-INs during extinction retrieval. Orange indicates No DBS manipulation during extinction training and green indicates 40 Hz DBS manipulation during extinction training. Data are mean ± SEM. \**P* < 0.05, unpaired Student’s t-test.

During extinction retrieval without DBS, only a fraction of PV-INs (50% of neurons with 0–15 Hz basal firing rate and 37.5% of neurons with 15–30 Hz basal firing rate) exhibited increased firing rates in response to the CS, depending on their basal firing rates (Figure 5, D and G). However, when paired with DBS, all three groups of PV-INs, including high-firing rate PV-INs, exhibited significant increases in firing rates in response to the CS (Figure 5, E, F and I). The firing frequencies of PV-INs shifted toward higher values (Figure 5, H and J) during CS presentation in the presence of DBS, with a shorter latency (Figure 5, K and L). In contrast, putative pyramidal neurons in the DBS group showed an inverse redistribution in firing rate changes, including a larger proportion with decreased firing rates during CS presentation (Extended Data Fig. 10), indicating increased inhibition. These results demonstrate that low-gamma DBS in the vCA1 region enhances the responsiveness of PV-INs, particularly those with higher basal firing rates, during extinction retrieval, while promoting inhibition of pyramidal neurons.

### Enduring activity of PV-INs by low-gamma DBS suppresses engram cells encoding fear memory in vCA1

Given that low-gamma vCA1 DBS enhanced PV-IN activity, we postulated that the robust suppression of DBS on cued fear responses could arise from its ability to inhibit fear engram cells. Using the targeted recombination in active populations (TRAP) strategy (41–43), we labeled fear engram cells by crossing FosCreERT2 (FosTRAP2) mice with the PV-Flp mouse line (FosTRAP2::PV-Flp). Co-administering Flp-dependent AAV-fDIO-GCaMP6m and Cre-dependent AAV-DIO-jRGECO1a into vCA1 allowed simultaneous monitoring of PV neuron and fear engram cell activities during extinction training paired with low-gamma DBS (Figure 6, A–C). Ca^2+^ signals revealed that PV neuron activity was significantly higher in the DBS group than in the no DBS group throughout extinction. In contrast, fear engram cells showed increased activity only during the Early- Ext. phase, inhibited by DBS, and decreased activity during the Late-Ext. phase, more pronounced with DBS (Figure 6, D and E).

**Figure 6.**
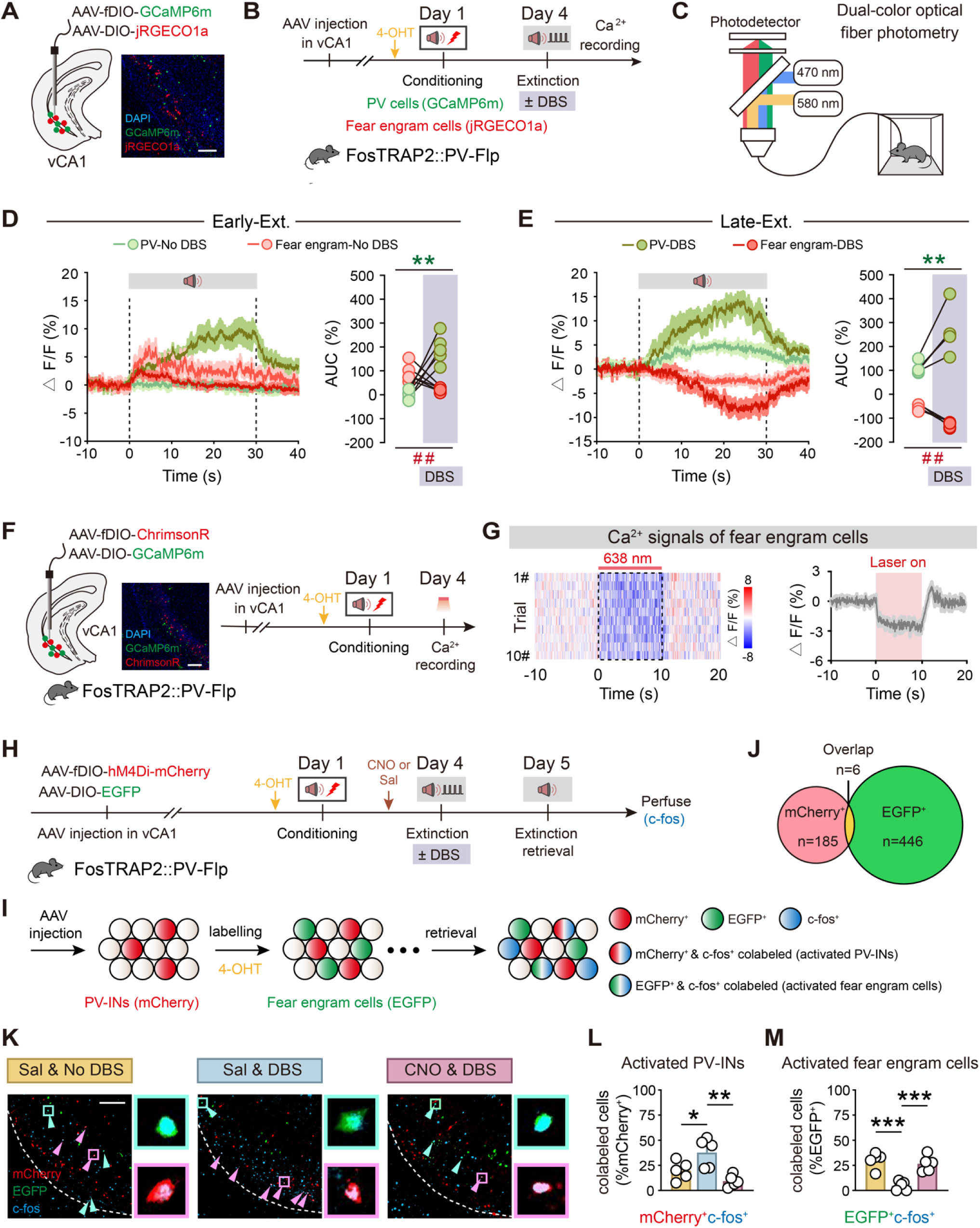
Low-gamma DBS paired extinction training engages vCA1 PV-INs to suppress fear engram cells. (**A**) Schematics of AAV injections and representative image of virus expression. Scale bar, 100 μm. (**B** and **C**) Schematics of experimental design. (**D** and **E**) Average calcium signals in PV-INs and fear engram cells during Early-Ext. (**D**) and Late-Ext (**E**). \*\**P* < 0.01, PV-INs-DBS *vs.* PV-INs-No DBS, paired Student’s t-test. ^##^*P* < 0.01, fear engram cells -DBS *vs.* fear engram cells-No DBS, paired Student’s t-test. DBS group, *n* = 5 mice, No DBS group, *n* = 5 mice. (**F**) Schematics of AAV injections and experimental design. Representative images of GCaMP6m expression in fear engram and ChrimsonR-expression in PV-INs in vCA1. Scale bar, 100 μm. (**G**) Representative heat map of fiber photometry recordings (left). Averaged fluorescence decreased in response to optogenetic stimulation (right) (*n* = 5 mice). (**H**) Schematics of AAV injections and experimental design. Administration of 4-OHT, 30 min before fear conditioning (i.p.), to FosTRAP2::PV-Flp mice was used to induce permanent expression of EGFP in neurons active around the time of the injection. (**I**) Genetic design to investigate fear engram cells and neurons activated during extinction retrieval. Red circles represent PV-INs, green circles represent neurons labeled during conditioning, and blue circles represent neurons activated during memory retrieval. (**J**) Overlap between vCA1 PV-INs (mCherry^+^) and fear engram cells (EGFP^+^). (**K**) Representative images of mCherry^+^ (red) and EGFP^+^ (green) and c-fos^+^ (blue) immunofluorescence in vCA1. Magenta arrowheads denote colabeled mCherry^+^/c-fos^+^ cells; cyan arrowheads denote colabeled EGFP^+^/c-fos^+^ cells. Circles represent enlarged images on the right. Scale bar, 100 μm. (**L** and **M**) The percentage of activated PV-INs (mCherry^+^/c-fos^+^) and activated fear engram cells (EGFP^+^/c-fos^+^). Data are mean ± SEM. \**P* < 0.05, \*\**P* < 0.01, \*\*\**P* < 0.001, unpaired Student’s t-test.

To validate the direct influence of PV-INs on fear engram cells, we introduced Flp-dependent AAV-fDIO-ChrimsonR and Cre-dependent AAV-DIO-GCaMP6m into vCA1 of FosTRAP2::PV-Flp mice. Activation of PV-INs through red light illumination in the vCA1 region significantly reduced fear engram cell Ca^2+^ signals (Figure 6, F and G). Subsequent c-fos analysis in these mice, injected with AAV-fDIO-hM4Di-mCherry and AAV-DIO-EGFP into vCA1, revealed that during extinction retrieval, the DBS group showed increased activated PV-INs (mCherry^+^/c-fos^+^) and decreased activated fear engram cells (EGFP^+^/c-fos^+^) compared to the no DBS group (Figure 6, H–M). Chemogenetic suppression of PV-INs prevented the DBS-induced reduction in fear engram cells, indicating that low-gamma vCA1 DBS activates PV-INs, leading to fear engram cell suppression and reduced cued fear responses (Figure 6, K–M). Importantly, there was minimal overlap between mCherry^+^ and EGFP^+^ cells (Figure 6, J and K), suggesting that the proportion of vCA1 PV-INs integrated into fear engram cells is negligible, and the PV-INs are preferentially engaged in fear extinction.

### Low-gamma DBS empowers LEC-vCA1 top-down feedforward inhibition pathway via PV-INs to suppress fear engram cells

To investigate the role of LEC-vCA1 pathway in mediating the effects of vCA1 DBS, we selectivity inhibited this pathway using chemogenetics during DBS. We found that inhibiting the LEC-vCA1 pathway (Figure 7, A–C), but not the MEC-vCA1 pathway (Supplemental Figure 11), attenuated the effects of vCA1 DBS, resulting in a higher fear response during extinction retrieval. Moreover, optically stimulating the LEC-vCA1 pathway with low-gamma frequency mimicked the effects of DBS on fear extinction (Supplemental Figure 12). This led us to hypothesize that DBS affects the inputs from LEC to vCA1 PV-INs, thereby suppressing fear engram cells. To test this, we introduced AAV-DIO-H2B-GFP into vCA1 and AAV-DIO-ChR2 into LEC of FosTRAP2::Sim1-Cre mice (Figure 7, D and E). Photostimulation of LEC fibers induced monosynaptic EPSCs and delayed inhibitory postsynaptic currents (IPSCs) in the same vCA1 fear engram cell, suggesting that LEC sends monosynaptic projections and a strong feedforward inhibitory circuit to fear engram cells. We found a significant increase in the amplitude of light-evoked IPSCs in vCA1 fear engram cells from DBS group, compared with those from the no DBS group, one day after fear extinction. The DBS group exhibited a marked increase in the IPSC/EPSC ratio (Figure 7, F and G). To confirm that vCA1 PV-INs are responsible for the feedforward inhibition within the LEC-vCA1 circuit, we blocked GABA release specifically from PV-positive interneurons via application of ω-agatoxin-IVA, a selective antagonist for P/Q-type calcium channels(44). After ω-agatoxin-IVA application, the IPSC amplitude exhibited a marked decrease (Figure 7, H and I), indicating that PV- INs mediate LEC SIM1^+^ layer 2a fan cells driven feed-forward inhibition onto vCA1 fear engram cells. Taken together, our findings suggest that low gamma DBS manipulation also strongly activates the inputs from LEC driving PV-INs-mediated feedforward inhibition in vCA1, leading to the long-term suppression of fear engram cells.

**Figure 7.**
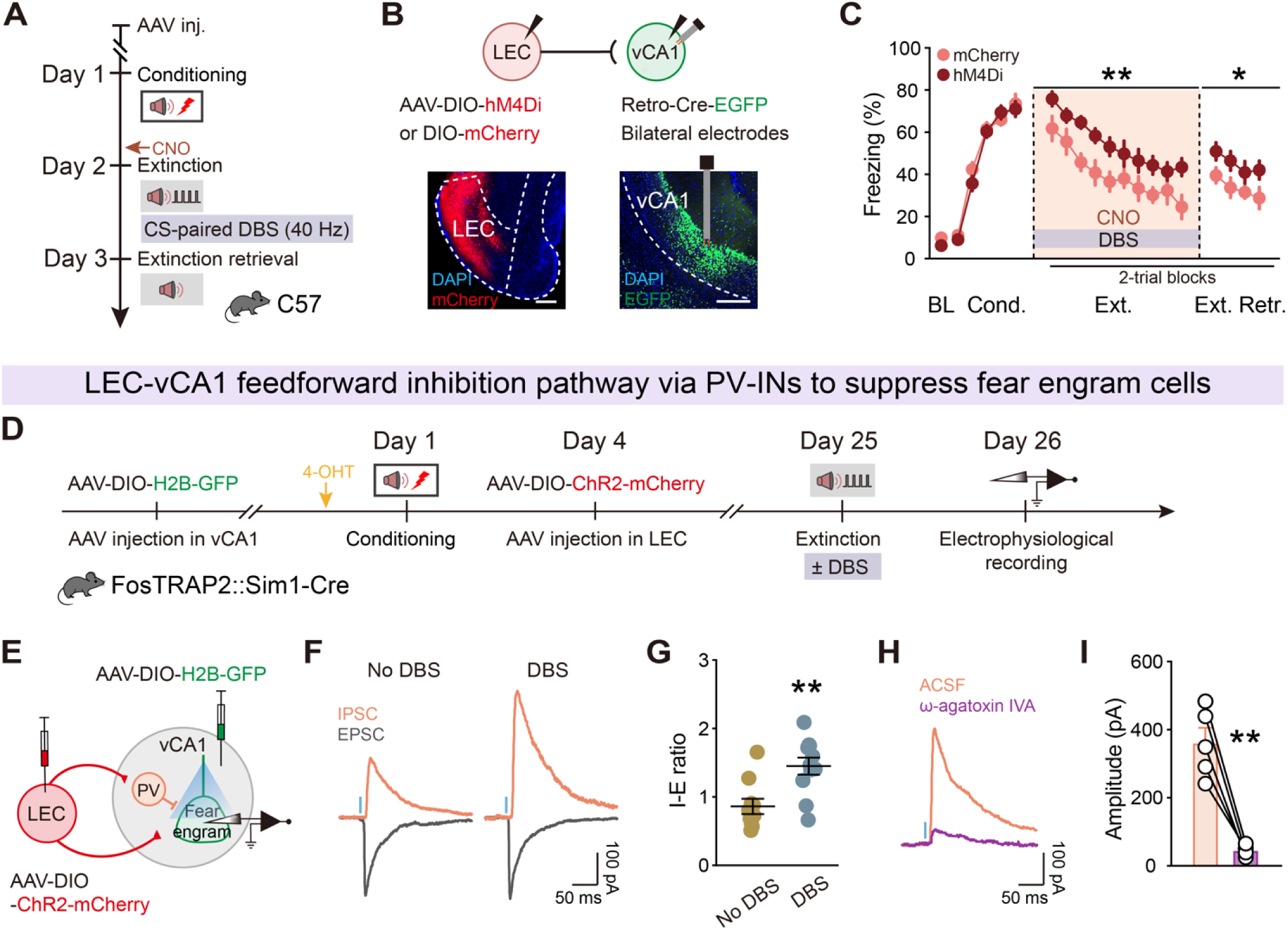
Low gamma DBS strengthens the inputs from LEC driving PV INs-mediated feedforward inhibition in vCA1 and induces long-lasting fear engram suppression. (**A**) Schematics of experimental design. CS is paired with 40 Hz DBS during extinction training and CNO was administrated 30 min (i.p.) before extinction training. (**B**) Schematics of AAV injections (top) and representative images of virus expression (bottom). Scale bar, 200 μm. (**C**) Effect of inhibiting LEC-vCA1 projectors on DBS-induced extinction promotion. Schematics of AAV injections. Time courses of freezing responses to the CS during fear conditioning, extinction training and extinction retrieval sessions. Statistics are as follows: two-way repeated measures ANOVA, main effect of AAV, conditioning, F_1,21_ = 0.4901, *P* = 0.4916; extinction training, F_1,21_ = 8.408, *P* = 0.0086; extinction retrieval, F_1,21_ = 7.556, *P* = 0.0120. mCherry group, *n* = 12 mice; hM4Di group, *n* = 11 mice. Data are mean ± SEM. \**P* < 0.05, \*\**P* < 0.01. (**D**) Schematics of AAV injections and experimental design. 4-OHT was administrated 30 min before fear conditioning. (**E**) Experimental scheme for simultaneous recording of light-evoked EPSCs and IPSCs on vCA1 fear engram cells. (**F**) Representative traces of EPSCs and IPSCs evoked by optogenetic stimulation of LEC fibers. (**G**) IPSC/EPSC peak ratios (No DBS, *n* = 10 cells; DBS, *n* = 11 cells). Data are mean ± SEM. \*\**P* < 0.01, unpaired Student’s t-test. (**H**) Representative traces showing that light-evoked IPSC amplitudes were reduced with application of 0.5 μM ω-agatoxin IVA. (**I**) Light-evoked IPSC amplitudes in vCA1 fear engram cells with and without ω-agatoxin IVA (*n* = 5 cells). Data are mean ± SEM. \*\**P* < 0.01, paired Student’s t-test.

### Low-gamma transcranial alternating current stimulation (tACS) targeting LEC enhances fear extinction

In light of the potential clinical applications of noninvasive neuromodulation, we explored the effects of transcranial alternating current stimulation (tACS) (38) on the LEC-vCA1 motif with the aim of utilizing electrical signals delivered via tACS to modulate the LEC, influencing vCA1 similarly to vCA1 DBS and ultimately promoting fear extinction. We implanted bilateral stimulation electrodes (anodes) over the LEC regions, while placing another stimulation electrode (cathode) on the neck skin of the mice. To evaluate the behavioral consequences of tACS, we divided the mice into two groups: those subjected to tACS (200 µA, 40 Hz, paired with the CS) and those without tACS (Figure 8, A and B). Compared to the No tACS group, the tACS group displayed accelerated extinction, and the effect persisted into the extinction retrieval session (Figure 8C).

**Figure 8.**
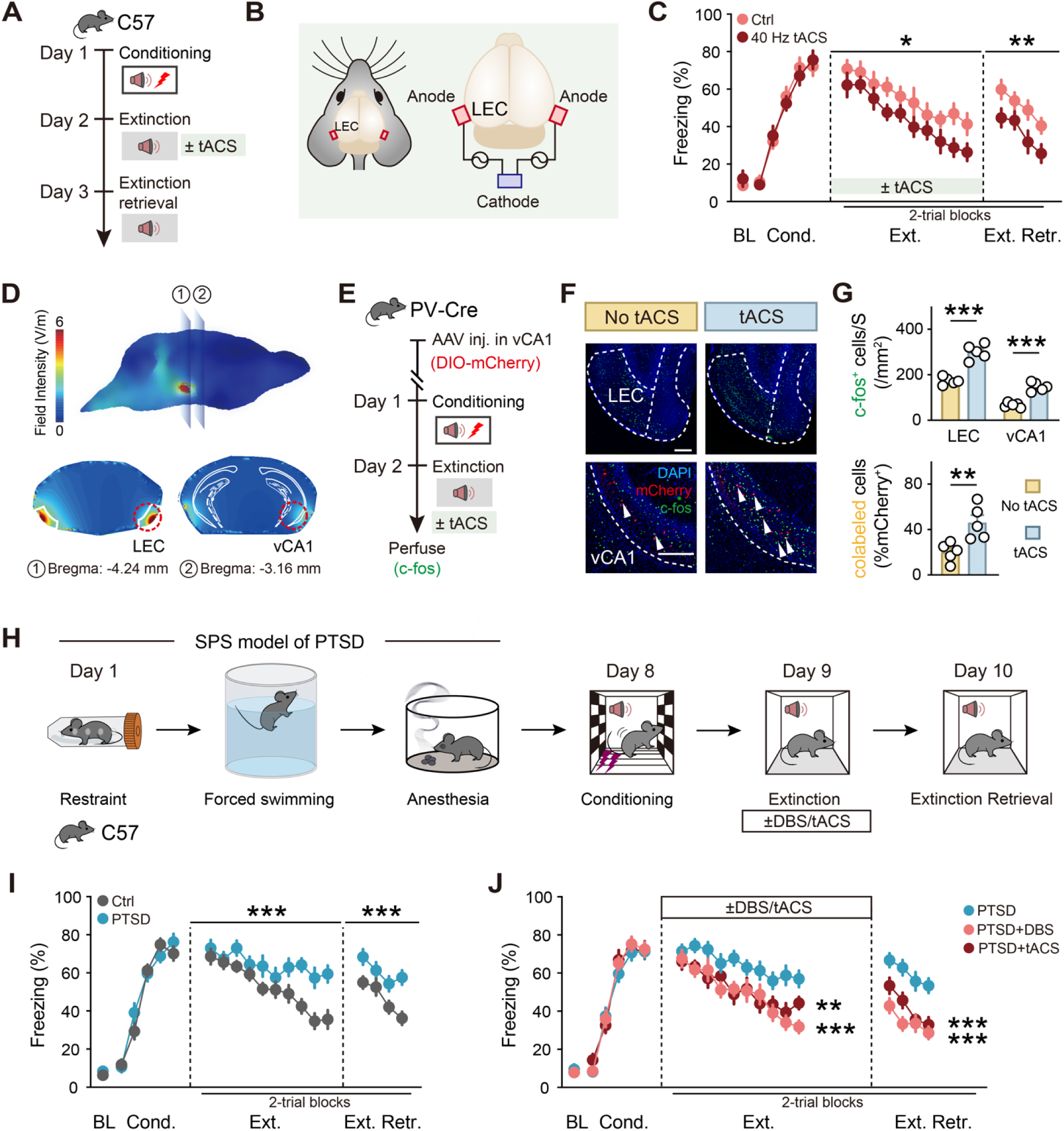
Low-gamma stimulation of LEC→vCA1 circuit removed fear memory, even under more traumatic conditions. (**A**) Schematics of experimental design. CS was paired with 40 Hz-tACS during extinction training. (**B**) Schematic diagram of stimulus configuration. Two stimulators were attached to the surface of LEC. The red electrodes were the anode and the blue electrode was the cathode. (**C**) Time courses of freezing responses to the CS during fear conditioning, extinction training and extinction retrieval sessions. Statistics are as follows: two-way repeated measures ANOVA, main effect of tACS, conditioning, F_1,14_ = 0.0331, *P* = 0.8582; extinction training, F_1,14_ = 8.055, *P* = 0.0132; extinction retrieval, F_1,14_ = 15.87, *P* = 0.0014. Ctrl group, *n* = 8 mice; 40 Hz tACS group, *n* = 8 mice. Data are mean ± SEM. \**P* < 0.05, \*\**P* < 0.01. (**D**) Predicted current density map at the surface of the brain during tACS (top) and slice images of the distribution showing peak current densities during tACS (bottom). (**E**) Schematics of experimental design. (**F**) Representative images of mCherry^+^ (red) and c-fos^+^ (green) immunofluorescence in the LEC and vCA1. White arrowheads denote colabeled cells. Scale bars, 200 μm. (**G**) Quantification for (**F**). Histograms represent mean ± SEM and circles denote individual mice. The number of activated (mCherry^+^/c-fos^+^) PV-INs was significantly higher in the tACS group than No tACS group. No tACS group, *n* = 5 mice; tACS group, *n* = 5 mice. N.S., no significant difference, \*\**P* < 0.01, \*\*\**P* < 0.001, unpaired Student’s t-test. (**H**) Schematic illustration of single-prolonged stress (SPS) and the fear-conditioning paradigm. CS was paired with 40 Hz DBS-vCA1 or tACS-LEC during extinction training. (**I**) SPS model of PTSD mice had impaired fear extinction. Time courses of freezing responses to the CS during fear conditioning, extinction training and extinction retrieval sessions (right). Statistics are as follows: two-way repeated measures ANOVA, main effect of AAV, conditioning, F_1,16_ = 0.2782, *P* = 0.6051; extinction training, F_1,16_ = 22.92, *P* = 0.0002; extinction retrieval, F_1,16_ = 38.08, *P* < 0.0001. Ctrl group, *n* = 9 mice; PTSD group, *n* = 9 mice. Data are mean ± SEM. N.S., no significant difference, \*\*\**P* < 0.001. (**J**) Effects of low-gamma stimulation of the LEC→vCA1 circuit on extinction training and extinction retrieval in SPS model of PTSD mice. Statistics are as follows: two-way repeated measures ANOVA. PTSD *vs.* PTSD+DBS, conditioning, F_1,16_ = 0.5860, *P* = 0.4551; extinction training, F_1,16_ = 16.79, *P* = 0.0008; extinction retrieval, F_1,16_ = 70.31, *P* < 0.0001. PTSD *vs.* PTSD+tACS, conditioning, F_1,15_ = 0.5624, *P* = 0.4649; extinction training, F_1,15_ = 14.42, *P* = 0.0018; extinction retrieval, F_1,15_ = 30.04, *P* < 0.0001. PTSD group, *n* = 9 mice, PTSD+DBS group, *n* = 9 mice, PTSD+tACS group, *n* = 8 mice. Data are mean ± SEM. N.S., no significant difference, \*\**P* < 0.01, \*\*\**P* < 0.001.

To further understand the potential impact of low-gamma (40 Hz) tACS on vCA1, we conducted computational modeling to assess the electric field generated during tACS. The predicted current density map at the brain surface, along with specific slice views, revealed an increase in the current density within vCA1 during tACS targeting LEC (Figure 8D). In parallel, we analyzed c-fos expression levels to quantify the activity patterns in the absence and presence of the 40 Hz tACS (Figure 8E). Compared to No tACS, the tACS group exhibited significant increases in the number of c-fos^+^ cells in both the LEC and vCA1 regions. Moreover, the number of PV^+^/c-fos^+^ cells in vCA1 was also substantially increased (Figure 8, F and G). These findings suggest that noninvasive LEC tACS shares similar cellular mechanisms with vCA1 DBS, primarily by recruiting vCA1 PV-INs to enhance neural communication between the cortex and hippocampus.

### Low-gamma LEC tACS and vCA1 DBS effectively reduce persistent fear in a mouse model of PTSD

Finally, given that conditions such as anxiety disorders and PTSD are characterized by the enduring nature of learned fear and are closely tied to deficiencies in extinction learning (1, 2), we assessed whether low-gamma LEC tACS or vCA1 DBS could alleviate fear in PTSD. We employed a mouse model of PTSD induced by single prolonged stress (SPS) (45–49), involving three consecutive stressors: restraint, forced swimming, and anesthesia (Figure 8H). Notoriously known for being resistant to fear extinction, this form of PTSD indeed exhibited obstinate fear memory, despite no discernible differences in the fear learning curve compared to the control mice (Figure 8I). Remarkably, the application of either LEC tACS or vCA1 DBS significantly enhanced the extinction of cued fear in the PTSD mice (Figure 8J). This finding underscores the effectiveness of these invasive and noninvasive neuromodulation approaches, which target low-gamma entrainment of entorhinal-hippocampal activity, in augmenting extinction processes, thereby facilitating the removal of traumatic memory, even under extreme conditions.

## Discussion

Gaining insights into the neurological mechanisms underlying fear extinction holds significant promise for psychotherapy, particularly for addressing the challenging issue of PTSD. Our current study unveils a direct projection pathway from the LEC to the vCA1, which is necessary and sufficient for implementing fear extinction. We unravel that fear extinction relies on low-gamma oscillations between the LEC and vCA1 at the circuit level coordinated by vCA1 PV-INs. Direct projections from LEC layer 2a fan cells to vCA1 PV-INs are distinct from indirect projections to the dorsal hippocampus. Furthermore, we found that exogenous low-gamma vCA1 DBS not only enhances fear extinction but also exerts enduring benefits. This remarkable efficacy is primarily attributed to the activation of high-firing PV-INs and the persistent suppression of fear engram cells, leading to a sustained reduction in fear responses. In our exploration of potential treatments for fear-related disorders like PTSD, we found that noninvasive low-gamma LEC tACS effectively reduces enduring fear when combined with fear extinction training. This positive outcome holds even in a mouse model of PTSD with the most extinction-resilient form of fear memory, rationalizing its practical utility. Together, our study uncovers an unknown top-down structural motif along the cortical-subcortical axis, in which interregional synchronization of low-gamma oscillations between LEC and vCA1 prompts extinction of enduring fear memory. These findings not only define the circuitry and mechanistic basis of fear extinction but also present a proof of principle for using FDA approved invasive and non-invasive approaches to hijack this novel pathway for removing traumatic memories with unprecedented efficacy and persistence (Figure 9).

**Figure 9.**
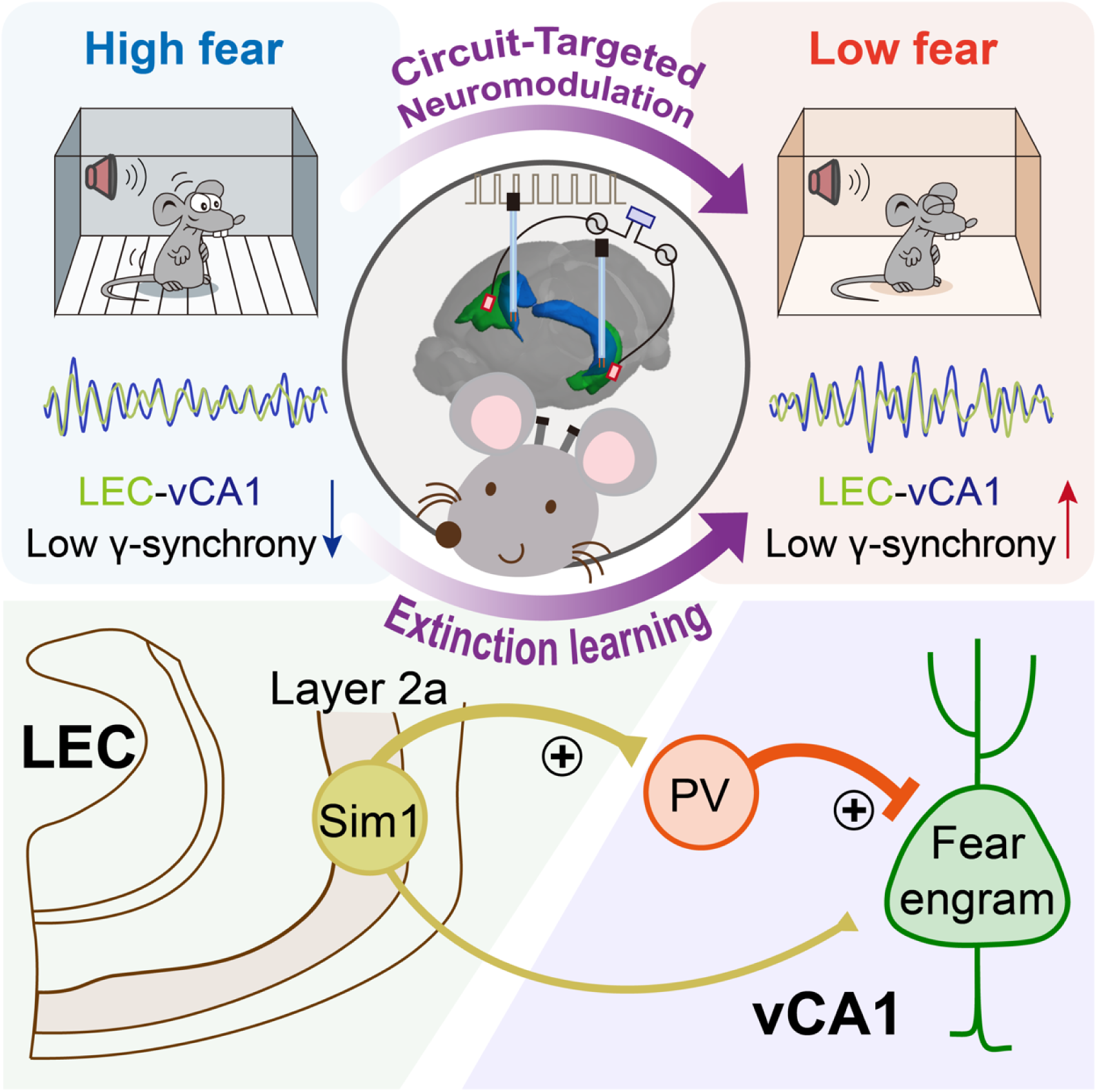
Scheme for a direct LEC-vCA1 projection pathway and the role of low-gamma oscillations and interregional entrainment in driving fear extinction, orchestrated by vCA1 PV-INs. This cortical-subcortical motif can be therapeutically targeted through either vCA1 DBS or LEC tACS to enhance feed-forward inhibition of fear engram neurons, thereby augmenting extinction to remove traumatic memories.

Contrary to the extensively studied dorsal hippocampal-entorhinal network, which supports both spatial navigation and associative memory (15–18, 50, 51), the connectivity, activity, and consequent behavioral implications of the ventral hippocampal-entorhinal network remain largely unexplored. A circuit mapping study has unveiled significant variations in input proportions and distributions between dorsal and ventral hippocampal CA1 pyramidal neurons, including distinct input patterns from the EC (52). Notably, there are instances where projections from EC neurons expressing corticotropin-releasing factor (CRF) directly target the vCA1, influencing behaviors of mice that respond to human experimenters’ sex and modulating the animals’ neural responses to ketamine (53). Here, we identify a direct projection from LEC layer 2a fan cells to vCA1 PV-INs, which controls fear extinction learning. Both populations of neurons in the projections, including LEC layer 2a fan cells and vCA1 PV-INs, are significantly activated by fear extinction learning. Notably, the LEC-vCA1 projections are necessary for the low-gamma-band oscillatory firing in each area, along with interregional low-gamma synchronization in response to fear extinction learning. Behaviorally, the specific activation or inhibition of this projection demonstrated a bidirectional influence on fear extinction. More strikingly, this projection was amenable to alteration through DBS and non-invasive tACS neuromodulation approaches. Among the projections from LEC layer 2a fan cells to various cell types in vCA1, isolation of this pivotal connection from LEC layer 2a fan cells to vCA1 PV-INs opens up an exciting avenue to decode the neural network mechanism of fear extinction. In conjunction with the well-known role of dorsal hippocampal-entorhinal circuits in spatial navigation and associative memory, our discovery of a previously unknown monosynaptic pathway from LEC to the vCA1 region, characterized by a unique projection pattern and specificity in fear extinction, exemplifies the organization of parallel structural and functional motifs for segregating single memory trace with diverse contents.

The LEC-vCA1 pathway, pivotal to fear extinction, is likely part of cognitive motif ensembles for memory processing. Unlike the established extinction circuits (22, 54–56) that collectively appear to affectively inhibit the expression of conditioned fear behaviors, the LEC integrates diverse sensory information (14, 18, 57–59), aided by dopaminergic innervation, to construct a cognitive map of abstract task rules (13). Fear extinction as a more abstract form of inhibitory learning (60) requires a dopaminergic switch for transitioning from fear to safety (61, 62). However, the contribution of LEC dopamine signals to the LEC-vCA1 motif remains open for future investigation. Thus, parallel cortical-subcortical motifs effectively process intricate contextual and sensory cues, with the LEC-vCA1 pathway being the key handle for implementing extinction of conditioned fear behaviors.

It is plausible that vCA1 PV-INs, within the top-down motif, are selectively and progressively recruited during fear extinction learning, facilitating synchronization of cortical and subcortical networks for fear extinction. This aligns with the concept that fear extinction involves inhibitory learning mechanisms directed against the original fear memory (63). In this study, we present compelling evidence supporting the existence of an extinction-initiated memory trace, with vCA1 PV- INs playing a causal role in suppressing fear memory engram cells at the network level. Notably, vCA1 PV-INs exhibit at least three forms of adaptations upon fear extinction. *Firstly*, neuronal activity increases gradually during fear extinction learning, exemplified by the progressive rise in cue-evoked Ca^2+^ signals. *Secondly*, there is a post-learning (extinction) adaptation, evident in elevated c-fos expression in vCA1 PV-INs following extinction learning compared to the control group in a homecage setting. *Lastly*, persistent plasticity of vCA1 PV-INs emerges during extinction retrieval, characterized by enhanced neuronal firing in response to CS presentation and a robust increase in the likelihood of high-firing rate PV-INs following vCA1 DBS modulation. While the precise molecular mechanisms of these vCA1 PV-IN adaptations are yet to be fully established, targeting such adaptations holds promise for translational applications in treating fear-related disorders. Presumably, vCA1 pyramidal neurons, as the final component of the cortical-subcortical motif for fear extinction, shift their firing towards low frequencies and more synchronous patterns due to vCA1 PV-IN adaptation to extinction training and vCA1 DBS. Overall, vCA1 PV-INs dynamically alter their activity throughout fear extinction processes, synchronizing neuronal activity within the cortical-subcortical motif for learning to extinguish fear memory.

Translating our circuit findings, we established two independent neuromodulation approaches, vCA1 DBS and LEC tACS, to enhance fear extinction, providing potential interventions for PTSD and other fear-related disorders. Both approaches effectively mitigated extinction resistance in a PTSD mouse model, attributed to activating high-firing rate vCA1 PV-INs and sustaining fear engram cell suppression, resulting in lasting fear reduction. Building on established therapeutic approaches for Parkinson’s disease using DBS (64, 65) and promising results in noninvasive brain stimulation methods, such as tACS, for various conditions (38, 66), our findings advocate applying neuromodulation techniques to address fear-related disorders, including PTSD. Because the neocortex is the most accessible with these neuromodulation technologies, our identification of adaptable motifs along the cortical-subcortical axis to boost fear extinction exemplifies the potential to advance the treatment options for individuals grappling with debilitating psychiatric and neurodegenerative conditions.

In conclusion, our study unveils the significance of the direct LEC-vCA1 projection and the role of low-gamma oscillations and interregional entrainment in driving fear extinction, orchestrated by vCA1 PV-INs. By successfully validating the efficacy of vCA1 DBS and LEC tACS, we introduce effective neuromodulation techniques to augment fear extinction, presenting promising interventions for PTSD and related disorders. These findings not only deepen our comprehension of psychotherapeutic approaches but also pave the way for innovative treatments in the realm of fear-related conditions. Our findings serve as a proof of principle for advancing therapies for memory diseases and neuropsychiatric disorders by precisely targeting accessible top-down cortical motifs in a pathway- and cell-specific manner.

## Methods

### Animals

The following animals were used in this study: C57BL/6J mice, Fos^2A-iCreER^ (TRAP2) (stock no. 030323) mice, PV-Cre (stock no.017320) mice, PV-Flp (stock no. 022730), SST-Cre (stock no. 013044) mice, VIP-Cre (stock no. 010908) mice, Sim1-Cre (stock no. 006395) mice and the lox-stop-lox-H2B- GFP (H2B-GFP^flox^) reporter mice. All mice were group-housed on a 12-h light/dark cycle with rodent chow and water *ad libitum*. Adult male mice (8–12 weeks old) were used for all experiments.

### Virus constructs

The following viruses were used: rAAV2/9-EF1α-DIO-eNpHR3.0-mCherry-WPRE-hGH polyA, rAAV-EF1α-DIO-RVG-WPRE-hGH polyA, rAAV-EF1α-DIO-H2B-EGFP-T2A-TVA-WPRE-hGH polyA, RV-EnvA-DG-DsRed, rAAV-nEf1α-fDIO-RVG-WPRE-hGH polyA, rAAV-nEF1α-fDIO- EGFP-T2A-TVA-WPRE-hGH polyA, rAAV2/9-EF1a-DIO-BFP-Flag-WPRE-bGH polyA, rAAV2/9-CaMKIIa-hChR2(E123T/T159C)-mCherry-WPRE-hGH polyA, rAAV2/9-CaMKIIa-mCherry- WPRE-hGH polyA, rAAV2/9-EF1α-DIO-hM4D(Gi)-mCherry-WPREs, rAAV2/9-EF1α-DIO-mCherry-WPRE-hGH polyA, rAAV2/9-EF1a-Con Fon-GCaMp6s-WPRE-hGH polyA, rAAV2/R- hSyn-FLP-WPRE-hGH polyA, rAAV2/9-EF1a-DIO-hM3D(Gq)-EGFP-WPREs, rAAV2/9-EF1a- DIO-EGFP-WPRE-hGH polyA, AAV2/1-hSyn-FLP-EGFP-WPRE-hGH polyA, rAAV2/9-hSyn-Con Fon-hM4D(Gi)-EGFP-WPRE-hGH polyA, rAAV2/9-hSyn-Con/Fon-EYFP-WPRE-hGH polyA, rAAV2/R-hSyn-CRE-EGFP-WPRE-hGH polyA, rAAV2/R-hSyn-CRE-WPRE-hGH polyA, rAAV2/9-nEF1a-fDIO-GCaMp6m-WPRE-hGH polyA, rAAV2/9-EF1a-jRGECO1a-WPREs, rAAV2/9-EF1α-DIO-hChR2(E123T/T159C)-mCherry-WPRE-hGH polyA and rAAV2/9-EF1a-DIO-EGFP-WPRE-hGH polyA (all purchased from Brain VTA, Wuhan, China); AAV2/9-hSyn-DIO- GCaMP6m-WPRE-pA, AAV2/9-hEF1a-fDIO-hM4D(Gi)-mCherry-ER2-WPRE-pA and AAV2/9-hEF1a-fDIO-ChrimsonR-mCherry-ER2-WPRE-pA (from Taitool Bioscience, Shanghai, China). All viruses were stored in aliquots at –80°C until use. The viral titers for injection were >10^12^ viral particles per ml.

### Stereotaxic surgery

Mice at 6–7 weeks old were anesthetized with 1% sodium pentobarbital via a single intraperitoneal injection per mouse (10 ml per kg body weight), after which each mouse was mounted in a stereotactic frame with non-rupture ear bars (RWD Life Science). After making an incision to the midline of the scalp, small craniotomies were performed using a microdrill with 0.5-mm burrs. For LFP recording, tungsten electrodes (Microprobes) were inserted into following coordinates (posterior to Bregma, AP; lateral to the midline, ML; below the Bregma, DV; in mm): LEC: AP, –4.16 mm; ML, ±3.90 mm; DV, –4.60 mm; MEC: AP, –4.46 mm; ML, ±3.0 mm; DV, –4.20 mm; vCA1: AP, –3.20 mm; ML, ±3.08 mm; DV, –4.0 mm.

For AAV injection, virus solutions were loaded into the tips of pipettes (Sutter Glass pipettes) and injected at the following coordinates: LEC: AP, –4.16 mm; ML, ±3.90 mm; DV, –4.60 mm; MEC: AP, –4.46 mm; ML, ±3.0 mm; DV, –4.20 mm; vCA1: AP, –3.20 mm; ML, ±3.08 mm; DV, –4.0 mm. After injection, the pipette was left in place for an additional 10 min to allow the injectant to diffuse adequately.

For retrograde monosynaptic tracing, a 1:1 volume mixture of AAV-EF1α-DIO-RVG and AAV- EF1α-DIO-H2B-EGFP-TVA (200-300 nl) was injected into the vCA1 of PV-Cre mice. A 1:1 volume mixture of AAV-EF1α-fDIO-RVG and AAV-EF1α-fDIO-H2B-EGFP-TVA (200-300 nl) was injected into the vCA1 of PV-Flp mice. Two weeks later, RV-ENVA-dG-DsRed (200-300 nl) was injected into the same location. The histology experiments were performed one week after rabies virus injection. For fiber photometry and optogenetic experiments, ceramic fiber optic cannulas (200 µm in diameter, 0.37 numerical aperture (NA), Hangzhou Newdoon Technology) were implanted above the vCA1 (AP, –3.20 mm; ML, ±3.08 mm; DV, –3.9 mm).

### Fear conditioning, extinction, and memory retrieval

All auditory fear conditioning, extinction, and memory retrieval procedures were performed using the Ugo Basile Fear Conditioning System (UGO BASILE srl). Briefly, mice were first handled and habituated to the conditioning chamber for three successive days. The conditioning chambers (17 cm × 17 cm × 25 cm) were equipped with stainless-steel shocking grids and connected to a precision- feedback current-regulated shocker. During fear conditioning, the chamber walls were covered with black-and-white checkered wallpaper, and the chambers were cleaned with 75% ethanol (context A). On day 1, mice were conditioned individually in context A with five pure tones (CS; 4 kHz, 76 dB, 30 s each) delivered at variable intervals (20-180 s). Each tone was co-terminated with a foot shock (US; 0.75 mA, 2 s each). ANY-maze software (Stoelting Co.) was used to automatically control the delivery of tones and foot shocks. Conditioned mice were returned to their homecages 30 s after the end of the last tone, and the cage was cleaned with 75% ethanol for each mouse. For extinction learning, mice conditioned on day 1 were presented with 20 CS presentations (4 kHz, 76 dB, 30 s each) without foot shock in context B (gray floor box) on day 2. On day 3, mice received eight CS-alone (30 s each) presentations in the extinction context (context B) for extinction retrieval. The movement of the mouse in the chamber was recorded using a near-infrared camera and analyzed in real-time with ANY-maze software. A fear response was operationally defined as measurable behavioral freezing (more than 1-s cessation of movement), which was automatically scored and analyzed by ANY-maze software.

### Open field test

The open field test was conducted to measure the effect of DBS or tACS on general locomotor activity. The mice with DBS or tACS electrodes were connected to the stimulator by means of a flexible cable and placed in an open field chamber (40-cm length, 40-cm width, 30-cm height). The open field chamber was divided into a center zone (center, 20 × 20 cm) and an outer zone (periphery). The movement of the mice was recorded and analyzed by using a video-tracking system (EthoVision 3.0, Noldus). The open field test consisted of an 18-minute session in which there were six alternating 3- minute epochs (OFF-ON epochs). The time spent in the center zone and the total distance and velocity traveled in the whole open field arena were measured over 18 min.

### Elevated plus maze test

The elevated plus maze was made of grey plastic and consisted of two closed arms (30 × 5 cm), two open arms (30 × 5 cm) and a central platform (5 × 5 cm). The maze was elevated 30 cm from the ground. The test mouse was placed in the center of the crossed maze, and the locomotion of the animal was recorded with a video-tracking system (EthoVision 3.0, Noldus) for 9 min. Each session was divided into three alternating 3-minute epochs: stimulation off, stimulation on, and stimulation off (OFF-ON epochs). The time spent in the open arms and that spent in the closed arms during the 9 mins period were quantified by EthoVision software.

### PTSD model

Single prolonged stress (SPS) was conducted as previously described(46). Briefly, the mice were restrained for 2 h in 50 mL polypropylene conical tubes with a nose hole for ventilation. Subsequently, the mice were forced to swim for 20 min in a plastic tub (20 cm diameter, 30 cm height) filled with water (24°C ± 1°C, 20 cm depth). Then, they were dried and allowed to recuperate for 15 min before being exposed to isoflurane and anesthetized until the characteristics of rapid breathing and loss of responses to toe and tail pinch were evident. After that, the mice were left undisturbed in their homecages for 7 days.

### LFP recording and analysis

Neural signals were recorded by using a multichannel data acquisition system (Zeus, Bio-Signal Technologies, Nanjing, China). Local field potentials were down-sampled to 1000 Hz prior to analysis and filtered with a bandpass filter between 0 and 150 Hz. Raw data were stored for later offline analysis. The time course of the LFP power spectra was generated using spectrogram analysis (NeuroExplorer, Nex Technologies) and the resultant three-dimensional time frequency spectra were smoothed using a Gaussian filter. To construct LFP power spectrograms, a multi-taper periodogram method was employed (5 tapers). Frequency-Power curve was generated using NeuroExplorer software Power Spectra for Continuous function.

Coherence between LFP channels across different electrodes was calculated using the weighted phase lag index (wPLI)(67). The wPLI is a measure of phase-synchronization between LFP signals, which is less affected by volume-conduction, noise and sample size. wPLI was estimated using the imaginary component of the (***S_xy_***) (Equations 1 and 2).

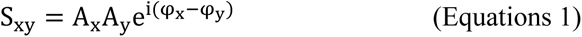

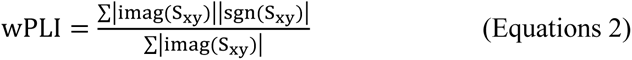

where ***A_x_*** and ***A_y_*** are instantaneous amplitudes; and ***φ_x_*** and ***φ_y_*** are instantaneous phases for vCA1 and EC (LEC or MEC) signals, respectively.

### Optogenetic manipulation

For optogenetic manipulation during behavioral assays, a 473-nm (blue light) or 589-nm laser (yellow light) (Hangzhou Newdoon Technology Co. Ltd) was connected to a patch cord with connectors on each end. For optogenetic inhibition of PV neurons, AAV-DIO-NpHR-mCherry or AAV-DIO- mCherry was injected into the vCA1 of PV-Cre mice, where optic fibers (200 μm in diameter, 0.37 NA) were implanted to allow inhibition at the cell bodies of PV neurons. To optogenetically inhibit LEC neuron terminals in the vCA1, AAV-DIO-NpHR-mCherry or control virus was injected in LEC and optical fibers were individually implanted into vCA1. The mice were tethered to optic fiber patch cords using ceramic mating sleeves and received photoinhibition (589 nm, 8-10 mW) in a continuous pattern during presentation of every 30-s CS (exceeding 5 s before and after the CS to ensure the light delivery covered the CS exposure).

For optogenetic stimulation of PV neurons and terminals from LEC and MEC with ChR2, mice were tethered to optic fiber patch cords and received photostimulation (473 nm, 4-6 mW) in 10 ms pulses at 40 Hz during presentation of every 30-s CS (exceeding 5 s before and after the CS to ensure the light delivery covered the CS exposure).

### Chemogenetic manipulation

For chemogenetic activation experiments, AAV-DIO-hM3Dq-EGFP or AAV-DIO-EGFP was bilaterally injected into LEC of Sim1-Cre mice and cannulas were implanted into vCA1. After four weeks to allow viral expression, clozapine N-oxide (CNO, 1 mM) was applied into the cannulas with a microinjection pump (RWD Ltd.) at a rate of 0.2 µL/min and then an additional minute was given to allow the drug to be locally delivered into the LEC axon terminal fields.

For LEC-vCA1 projection inhibition, AAV2/1-Flp was injected into LEC and AAV-Cre (on)/Flp (on)-hM4Di-EGFP or AAV-Cre (on)/Flp (on)-EGFP into vCA1 of PV-Cre mice. For chemogenetic inhibition experiments, AAV-DIO-hM4Di-mCherry or AAV-DIO-mCherry was injected into vCA1 of PV-Cre, SST-Cre or VIP-Cre mice. After four weeks, mice were subjected to fear learning and extinction. CNO (1 mg/kg) was injected intraperitoneally 30 min before extinction training.

To inhibit vCA1-projecting LEC or MEC neurons, Retro-Cre-EGFP was injected into vCA1 and AAV-DIO-hM4Di-mCherry or control virus into LEC and MEC, respectively. After four weeks, mice were subjected to fear learning and extinction. CNO (1 mg/kg) was injected intraperitoneally 30 min before extinction training.

### Fiber photometry

The fiber photometry system (ThinkerTech, Nanjing, China) allows for real-time recording of fluorescence signals from genetically encoded calcium indicators in freely moving mice. AAV-DIO- GCaMP6m was injected and optical fibers (200 μm in diameter, 0.37 NA) were individually implanted into vCA1 in PV-Cre, SST-Cre or VIP-Cre mice. To record the calcium signals of vCA1-projecting LEC SIM1^+^ layer 2a fan cells, AAV-Cre(on)/Flp(on)-GCaMP6s and Retro-Flp were individually injected into LEC and vCA1 in Sim1-Cre mice. Optical fibers were implanted into LEC. To record the effect of DBS manipulation (different frequencies) on calcium signals in PV neurons, AAV virus expressing GCaMP6m was injected and optical fibers were implanted in PV-Cre mice.

To record calcium signals in fear engram cells induced by optogenetic stimulation of vCA1 PV-INs, FosTRAP2::PV-Flp mice were injected with AAV-DIO-GCaMP6m and AAV-DIO-ChrimsonR at vCA1 for expression in fear engram cells and PV neurons, respectively. Recordings in fear engram cells were performed concurrently with PV neurons being stimulated through the same optic fiber. Optogenetic stimulation was delivered (λ = 638 nm) at 40 Hz and fiber photometry recordings were made using an Arduino board running custom code synchronized to the photometry system.

To simultaneously record calcium signals of PV neurons and fear engrams in vCA1 at a 100-Hz sampling rate, different excitation wavelengths were used (470 nm for Ca^2+^-dependent GCaMP6m activity, 580 nm for Ca^2+^-dependent jRGECO1a activity). To minimize photo-bleaching, the laser power at the tip of the optical fiber was adjusted to a low level of 40-60 µW.

The emitted fluorescence signals were collected and converted into electrical signals to reflect neural activity. The analog voltage signals were digitalized at 100 Hz using a Power 1401 digitizer and Spike2 software (CED, Cambridge, UK). Photometry data were further analyzed using MATLAB. The data were segmented and aligned to the onset of trigger events within individual trials. Fluorescence changes (*ΔF/F*) were calculated as (*F – F_0_*)*/F_0_*, where *F_0_* is the mean fluorescence signal for 10 s before the trigger event. *ΔF/F* values are presented as heatmaps, single trial curve or average plots with the shaded area indicating the SEM.

### Deep brain stimulation (DBS)

The implanted bipolar DBS electrode made of platinum-iridium wire (76.2 μm diameter, coated, AM Systems) was connected to the stimulator by means of a flexible cable, allowing free movement of the animal. Mice were housed individually after the surgery during the entire period of the experiments.

Electrical stimulation was delivered through a stimulator (STG4008; Multi Channel Systems, Reutlingen, German), which generates square-wave biphasic current pulses. Mice received stimulation at 20 Hz (50 μA and pulse width of 5 ms), 40 Hz (50 μA and pulse width of 5 ms) or 130 Hz (50 μA and pulse width of 70 μs). In sham-DBS control experiments, mice were connected to the external cable but not subjected to electrical stimulation. Current and frequencies were chosen based on previous studies in the literature (68).

### Transcranial alternating current stimulation (tACS)

Upon exposure of the skull, a burr hole craniotomy was drilled using drill bit in the skull at a location chosen to target LEC (AP, –4.16 mm; ML, ±4.20 mm; DV, –4.85 mm). The dura was left intact. Bone anchor screws were driven approximately half-way through the skull and served as anode electrodes for tACS. Another electrode was inserted into the neck muscles and served as cathode electrodes. The stimulation group received tACS with 40 Hz of 200 μA over the bilateral LEC (STG4008; Multi Channel Systems, Reutlingen, German).

### Computational model and solution method

The Finite Element Method (FEM) was used to simulate the electric fields engendered by transcranial electric stimulation. The FEM model was meticulously constructed based upon the Digimouse dataset (69), a comprehensive whole-body computed tomography compendium of a mouse, boasting a high resolution (∼0.1 mm), curated by Alekseichuk et al. (70).

In the initial phase of the current study, segmentation of brain tissues was executed, and integration of the electrode was accomplished utilizing the ITK-SNAP software (71). The placement of all electrodes was meticulously aligned with the protocols used in the animal experiments. Subsequently, a tetrahedral-based FEM model was made to incorporate in excess of 5 million tetrahedral elements, through a series of modified segmentation procedures and the application of iso2mesh (72). Then, the FEM model was processed using Gmsh (73) and subsequently exported as a Nastran Bulk Data File for further computational analysis.

To define conductivities and compute the physics equations, COMSOL Multiphysics 4.3 (COMSOL, Inc., Burlington, MA) was employed. The conductivities were defined as follows (in Siemens per meter, S/m): white matter and grey matter, 0.126; Cerebrospinal Fluid (CSF), 1.654; Bone, 0.01; Scalp, 0.465; Eye Balls, 0.5; Copper, 7×10^6^; Platinum-iridium alloy, 6×10^7^.

The nature of the study was delineated as steady-state current studies, and the initial state of the electric field was predetermined to be zero. The electric field was derived through the injection of current (200 μA) via a stimulator (STG4008; Multi Channel Systems, Reutlingen, German). The magnitude of the electric field (*E*) was calculated as follows (Equations 3):

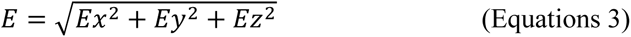

where *x*, *y*, and *z* represent the respective orthogonal directions.

### Histology and fluorescent immunostaining

Animals were deeply anesthetized with 1% sodium pentobarbital and were transcardially perfused with 1 × PBS, followed by 4% paraformaldehyde (PFA) in 1 × PBS. Brains were extracted and post-fixed overnight at 4°C in 4% PFA (pH = 7.2) and 40-μm coronal sections were collected using a vibratome (VT1000S, Leica). After a 15-min incubation in a 4,6-Diamidino-2-phenylindole dihydrochloride hydrate (DAPI) solution (1: 2000), sections were washed three times (15 min each time) in PBS with 0.1% Tween-20. Slides were mounted in the dark with glass coverslips using mounting media. The coverslips were sealed to the slide with nail polish. All fluorescent images were collected by taking serial z-stack images through 10 × or 20 × objectives of a confocal microscope (Digital Eclipse A1R+, Nikon).

For c-fos staining, brain slices were washed three times (5 min each time) with 1 × PBS and then blocked with 10% normal donkey serum in 1 × PBS with 0.3% Triton X-100 (PBST) for 1 h, after which they were incubated overnight at 4°C with rabbit anti-c-fos (1:500, Cell Signaling Technology, catalog no. 2250S). Sections were then washed with PBS, incubated in 2% normal donkey serum for 10 min, and then incubated for 2 h with Alexa Fluor 488 donkey anti-rabbit IgG or Alexa Fluor 568 donkey anti-rabbit (ThermoFisher Scientific).

For labeling PV, SST, VIP or Reelin, mice were perfused under deep anesthesia with PBS followed by 4% PFA. The brains were post-fixed in PFA for 2 h and stored in a 30% sucrose solution for 1-2 days at 4 °C. Brain sections (40-μm) were prepared with a cryostat (CM1900, Leica). Slices were incubated in the blocking solution with mouse anti-PV (1:300, Sigma-Aldrich, catalog no. MAB1572), goat anti-somatostatin (SST) (1:500, Santa Cruz, catalog no. sc-7819), rabbit anti-VIP (1:500, Immunostar, catalog no. 20077) or rabbit anti-Reelin (1:500, Invitrogen, catalog no. PA5-78413) at 4°C overnight and then with the secondary antibody (Alexa Fluor 568 donkey anti-mouse IgG, Alexa Fluor 488 donkey anti-mouse IgG, Alexa Fluor 647 donkey anti-mouse IgG, Alexa Fluor 568 donkey anti-goat IgG, Alexa Fluor 568 donkey anti-rabbit IgG or Alexa Fluor 647 donkey anti-rabbit IgG (all 1:500, Thermo Fisher Scientific) for 2 h. Slices were washed in 1 × PBS with 0.1% Tween-20, mounted onto slides, and covered with coverslips with ProLong Gold Antifade Mountant (Invitrogen).

### Slice electrophysiology

Whole-cell recordings were performed in acute brain slices. Mice were deeply anesthetized with 1% sodium pentobarbital and subsequently decapitated. Brains were dissected quickly and chilled in well- oxygenated (95% O_2_/5% CO_2_, v/v) ice-cold artificial cerebrospinal fluid (ACSF) containing the following (in mM): 125 NaCl, 2.5 KCl, 12.5 D-glucose, 1 MgCl_2_, 2 CaCl_2_, 1.25 NaH_2_PO_4_, and 25 NaHCO_3_ (pH 7.35-7.45). Coronal brain slices (300-µm thick) containing regions of interest were cut with a vibratome (Leica VT1000S, Germany). After recovery for 1 h in oxygenated ACSF at 30 ± 1°C, each slice was transferred to a recording chamber and was continuously superfused with oxygenated ACSF at a rate of 1–2 ml per minute. The neurons in vCA1 were patched under visual guidance using infrared differential-interference contrast microscopy (BX51WI, Olympus) and an optiMOS camera (QImaging). The slices were continuously perfused with well-oxygenated ACSF at 35 ± 1°C during all electrophysiological studies. Whole-cell patch clamp recordings were performed using an Axon 200B amplifier (Molecular Devices). Membranous currents were sampled and analyzed using a Digidata 1440 interface and a personal computer running Clampex and Clampfit software (Version 10.5, Axon Instruments). Optical stimulation of ChR2-, or NpHR-expressing neurons was performed using a collimated LED (Lumen Dynamics) with peak wavelengths of 473, 589 nm, respectively. The LED was connected to an Axon 200B amplifier to trigger photostimulation. The brain slice in the recording chamber was illuminated through a 40 × water-immersion objective lens (LUMPLFLN 40XW, Olympus). To verify the functional potency of NpHR-mediated optogenetic inhibition, yellow light (λ = 589 nm, 1-s pulse) was delivered to generate outward photocurrents under voltage-clamp mode, which promoted membrane hyperpolarization to reduce spikes produced by current injection under current-clamp mode. The functional potency of the ChR2-expressing virus was validated by measuring the number of action potentials evoked by using 40 Hz of blue-light stimulation (1 ms, λ = 473 nm). To evoke synaptic responses of PV neurons in the vCA1 by optogenetic photostimulation of LEC axons, the slice was illuminated with blue-light pulses of 5-ms durations. For light-evoked EPSCs, the recording pipettes (3–5 MΩ) were filled with a solution containing the following (in mM): 132.5 cesium gluconate, 17.5 CsCl, 2 MgCl_2_, 0.5 EGTA, 10 HEPES, 4 Mg-ATP, and 5 QX-314 chloride (280–300 mOsm, pH 7.2 with CsOH). TTX (1 μM TTX), 4-AP (100 μM), and NBQX (10 μM) were diluted in ACSF and applied through superfusion. For the feedforward inhibition test, light-evoked EPSCs and IPSCs were recorded at −70 mV and 0 mV, respectively, with 5-ms blue laser pulses every 20 s. For pharmacological isolation of PV-IPSCs specifically, PV-IPSCs were blocked using 0.5 μM of ω-agatoxin IVA, a selective antagonist for P/Q-type calcium channels (ApexBio, B7192). Data were analyzed by the Mini Analysis Program (Synaptosoft) with an amplitude threshold of 20 mV.

### Engram labeling

Recombination was induced with 4-hydroxytamoxifen (4-OHT, Sigma-Aldrich, Catalog no. H6278). In brief, 4-OHT was dissolved at 20 mg/ml in ethanol by shaking at 37°C for 30 min and then stored in aliquots at –20°C for up to several weeks. Before use, 4-OHT was re-dissolved in ethanol by shaking at 37°C. Corn oil (Sigma-Aldrich, Catalog no. C8267) was then added for a final concentration of 10 mg/ml 4-OHT, and the ethanol was evaporated by vacuum under centrifugation. The final 10 mg/ml 4-OHT solutions were stored at 4°C for < 24 h before use. All injections were delivered intraperitoneally (i.p.). Mice were transported from the vivarium to an adjacent holding room at least 3 h before the 4-OHT injection to minimize transportation-induced immediate early gene activity. Activity-dependent neuronal labeling was induced by a single intraperitoneal injection of 4-OHT (50 mg/kg mice) administered before the conditioning session for the fear-labeled mice. Mice were then returned to the vivarium with a regular 12 h light-dark cycle for the remainder of the experiment.

### *In vivo* electrophysiological recording

For in vivo optogenetic tagging of PV-INs, PV-Cre mice were unilaterally injected with AAV DIO ChR2-mCherry at vCA1 (AP, –3.20 mm; ML, ±3.08 mm; DV, –4.0 mm). Three weeks later, custom- made optrodes (consisting of an optical fiber and 8 tetrodes) were implanted at the same coordinates where the virus was injected. The tip of the optic fiber was 300 μm above the tip of the electrodes. Each tetrode was made of four twisted fine platinum/iridium wires (12.5 μm diameter, California Fine Wire). Silver wires with two screws were attached to the skull as ground. After a 7-day recovery from the surgery, the mice were habituated to the headstage and cables connected to the electrode on their heads for several days prior to electrophysiological recordings. Spiking activities were digitized at 30 kHz, band-pass filtered between 250 and 8000 Hz, and stored on a PC for further offline analysis.

### Spike sorting and unit classification with optogenetic tagging

Briefly, data were exported to Offline Sorter (Plexon Inc.) and NeuroExplorer (Nex Technologies) for offline analysis. Spike waveforms were identified by threshold crossing and sorted into units (presumptive neurons) by principal component analysis (PCA). Waveforms with inter-spike intervals (ISIs) less than the refractory period (2 ms) were excluded. Only units showing spikes with a signal- to-noise ratio larger than 2 were considered as neurons and included. Cross-correlation histograms were plotted to ensure that no unit was discriminated against more than once on different channels. All units were further classified as wide-spiking (WS) putative pyramidal neurons, narrow-spiking (NS) INs and fasts-piking parvalbumin (FS-PV) INs based on the peak-trough latency and the baseline firing rate. A unit with > 400 µs of peak-trough latency was classified as a WS neuron, with ≤400 µs of peak-trough latency was classified as an NS-IN and NS-IN with firing rate > 10 Hz was classified as an FS-PV IN.

For optical identification of vCA1 PV neurons, blue-light pulses (470 nm, 5-ms pulse duration, 1 Hz) were delivered at the end of each recording session. To assess whether these units were driven directly by ChR2 or indirectly by synaptic connections, we analyzed the onset latency relative to each light pulse. Only spikes with short spike latency (< 5 ms) and low jitter (< 3 ms) after light-pulse illumination were considered as being directly stimulated in this study. Only when the waveforms of laser-evoked and spontaneously generated spikes were highly similar (correlation coefficient >0.9), then they were considered to originate from the same unit.(74)

### Single unit firing analysis

To analyze vCA1 neuronal activity during 30-s trials, z-scored peristimulus time histograms (PSTHs) were calculated for each individual neuron, averaged over 8 CS trials for extinction retrieval, with or without DBS-vCA1 in extinction learning. Spikes were divided into 500-ms bins for visualizing individual unit responses and comparing responses between groups. A z-score was calculated for each bin relative to the 10-s prestimulus activity by subtracting average firing rate during baseline and by dividing the difference by the baseline standard deviation. For a given unit, firing rates during extinction retrieval and during baseline were compared to determine the significance of firing rate difference between these two conditions (paired t test). To compare PSTHs between groups, the mean z-score was calculated for each unit by averaging z-scored PSTHs during extinction retrieval (30-s CS). Response heatmaps were generated from z-scored PSTHs of units sorted by their mean z-score during 30-s CS in extinction retrieval.

### Quantification and statistical analysis

Detailed statistical analyses were performed using MATLAB and GraphPad Prism. The data were collected and processed randomly. All behavioral tests and analyses were blindly conducted. Data distributions were tested for normality and variance equality among groups was assessed using Levene’s test. Data are mean ± the standard error of the mean (S.E.M.) unless indicated otherwise. Statistical comparisons were performed using two-tailed unpaired Student’s *t* tests or two-tailed paired Student’s *t* tests, as well as one-way analyses of variance (ANOVAs) or two-way repeated-measures ANOVAs, where appropriate. For non-parametric datasets, Wilcoxon signed-rank test was used to determine significance. For *post-hoc* analysis, Bonferroni’s corrections or Sidak’s multiple comparisons test was used for multiple comparisons. Significance is mainly displayed as \**P* < 0.05, \*\**P* < 0.01, \*\*\**P* < 0.001; N.S. denotes no significant difference, which is not typically indicated except for emphasis.

### Study approval

All animal care and experimental procedures were approved by the Animal Ethics Committee of Shanghai Jiao Tong University School of Medicine and by the Institutional Animal Care and Use Committee (Department of Laboratory Animal Science, Shanghai Jiao Tong University School of Medicine; Policy Number DLAS-MP-ANIM. 01–05).

### Data and code availability

All data needed to evaluate the conclusions of the present study are present in the main paper and/or the supplemental information. This paper did not produce original code. Any additional information required for reanalyzing the reported data is available from the lead contact upon request.

## Acknowledgements

We thank Profs. Yi Wang (Zhejiang Chinese Medical University) and Jiamin Xu (East China Normal University) for their technique assistance. We are grateful to Profs. Xiaoming Li (Zhejiang University), Jiangteng Lv (Shanghai Jiao Tong University), Ju Huang (Shanghai Jiao Tong University), and Kexin Yuan (Tsinghua University) for providing transgenic mice used in this study in a generous manner. We thank Prof. Juan Song at University of North Carolina at Chapel Hill for critical comments on the manuscript. This study was supported by grants from the STI2030-Major Projects (2021ZD0202800), the National Natural Science Foundation of China (31930050, 81961128024, 32071023, 32200821, 32371078, and 32300843), the Science and Technology Commission of Shanghai Municipality (22XD1420700 and 23YF1433900), the Shanghai Municipal Health Commission (2022XD046), and innovative research team of high-level local universities in Shanghai.

## Author contributions

Z.-J.L., X.G., T.-F.Y., W.-G.L., and T.-L.X. conceived the project, designed the experiments, and interpreted the results. Z.-J.L. and X.G. performed the majority of behavioral experiments, animal surgery, immunohistochemistry, and data analysis. Y.-J.W., Q.W., and X.-Y.Z. assisted with some of the behavioral experiments and conducted viral injections. W.-K.G. assisted with DBS and tACS experiments. M.W. and Q.L. did computational modeling. X.G., Y.-J.W., and X.-R.W. performed slice recording and data analysis. Z.-J.L., X.G., M.X.Z., L.-Y.W., W.-G.L. and T.-L.X. wrote the manuscript with contributions from all authors. All authors read and approved the final manuscript.

## Additional information

Supplemental Material includes 13 figures and their legends are available for this paper online.

## Conflict of interest

The authors have declared that no conflict of interest exists.

## Supplemental Figures and Legends

The file includes:

Supplemental Figures 1 to 13

**Figure S1.**
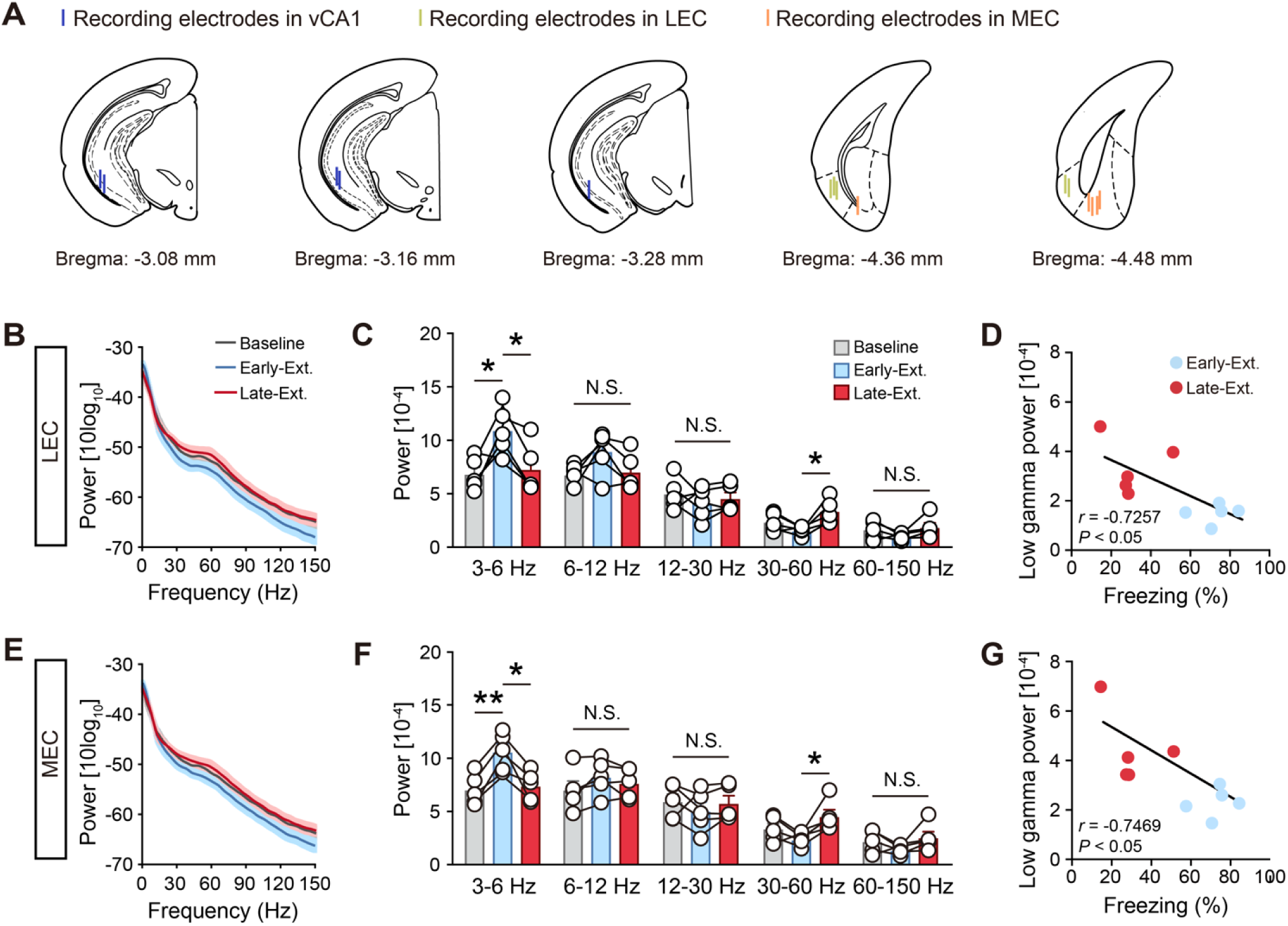
Fear extinction recruits low-gamma oscillations in the LEC and MEC. (**A**) Location of center for LFP electrode lesions for all mice. (**B** and **E**) Power spectrum of LEC (**B**) and MEC (**E**) LFP recordings during Baseline, Early-Ext. and Late Ext.. Solid lines represent the averages and shaded areas indicate SEM. (**C** and **F**) Average power of LEC (**C**) and MEC (**F**) LFP recordings during Baseline, Early-Ext. and Late Ext.. Histograms represent mean ± SEM and circles denote individual mice. *n* = 5. N.S., no significant difference, \**P* < 0.05, paired Student’s t-test. (**D** and **G**) Linear regression of freezing responses vs LEC (**D**) and MEC (**G**) low-gamma power during Early-Ext. and Late Ext. sessions.

**Figure S2.**
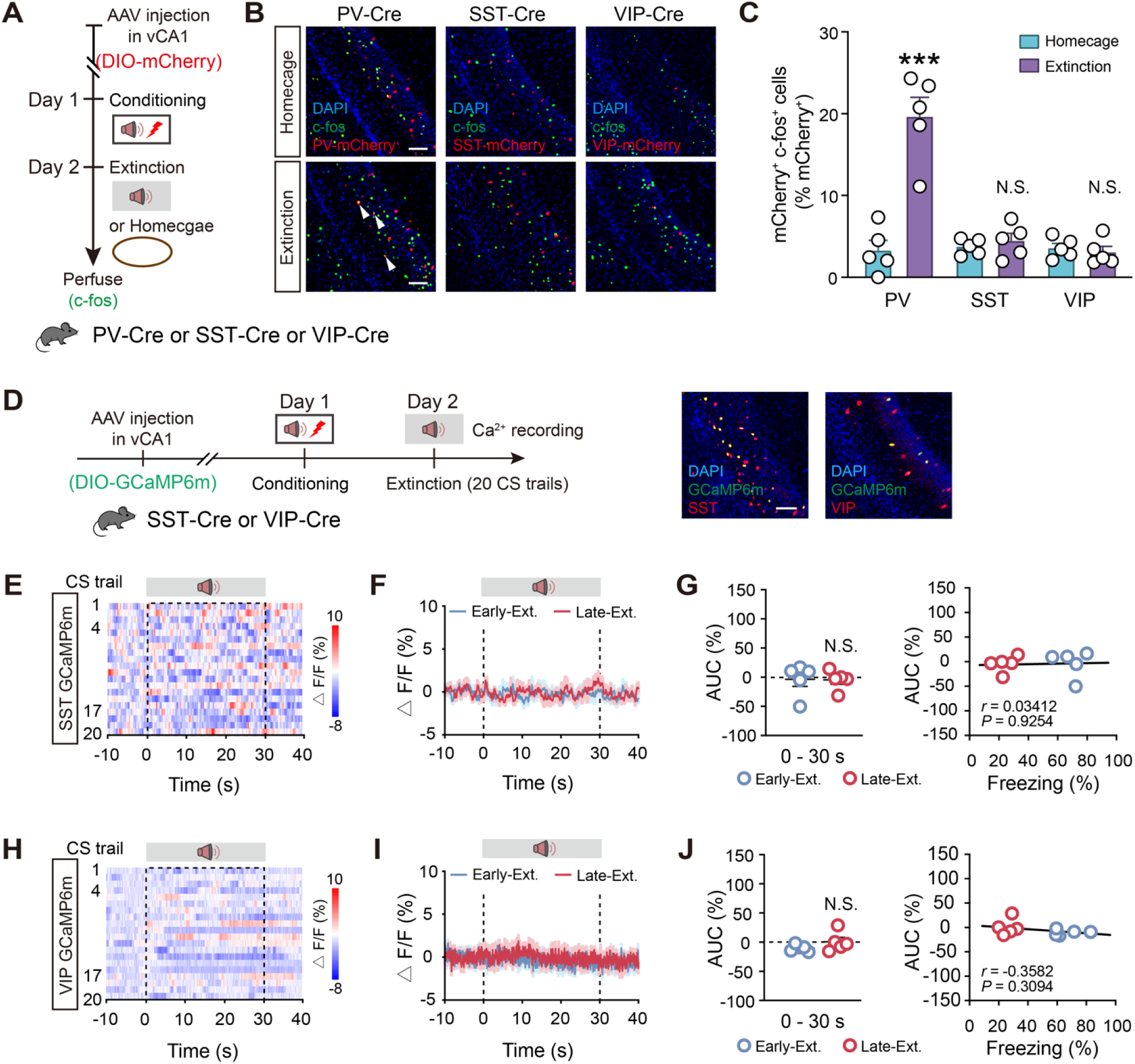
vCA1 PV-INs were activated in the extinction. (**A**) Schematics of AAV injections and experimental design. (**B**) Representative images of mCherry^+^ (red) and c-fos^+^ (green) immunofluorescence in the vCA1. The white arrowheads denote colabeled mCherry^+^/c-fos^+^ cells. Scale bar, 100 μm. (**C**) Histograms represent mean ± SEM and circles denote individual mice. The number of reactivated (mCherry^+^/c-fos^+^) PV-INs was significantly higher in the extinction group than the homecage group. Extinction group, *n* = 5 mice; homecage group, *n* = 5 mice. N.S., no significant difference, \*\*\**P* < 0.001, unpaired Student’s t-test. (**D**) Schematics of AAV injections, experimental design and immunostaining confirming the specificity of GCaMP6m expression in the SST-INs and VIP-INs. Scale bar, 100 μm. (**E**) Heatmap of calcium signals in the SST-INs during extinction training sessions. (**F**) Average calcium signals in the SST-INs during Early-Ext. and Late-Ext. Data are mean ± SEM. *n* = 5 mice. (**G**) Activity of the SST-INs and correlation of freezing responses with the calcium signals during Early-Ext. and Late-Ext. Quantification of AUC of calcium signals in the SST-INs (left). Data are mean ± SEM. N.S., no significant difference, paired Student’s t-test. Linear regression of freezing responses vs AUC of SST-INs GCaMP signals during Early-Ext. and Late Ext. (right). (**H**–**J**) The same as (**E**–**G**) for the VIP-INs. Data are mean ± SEM. *n* = 5 mice. N.S., no significant difference.

**Figure S3.**
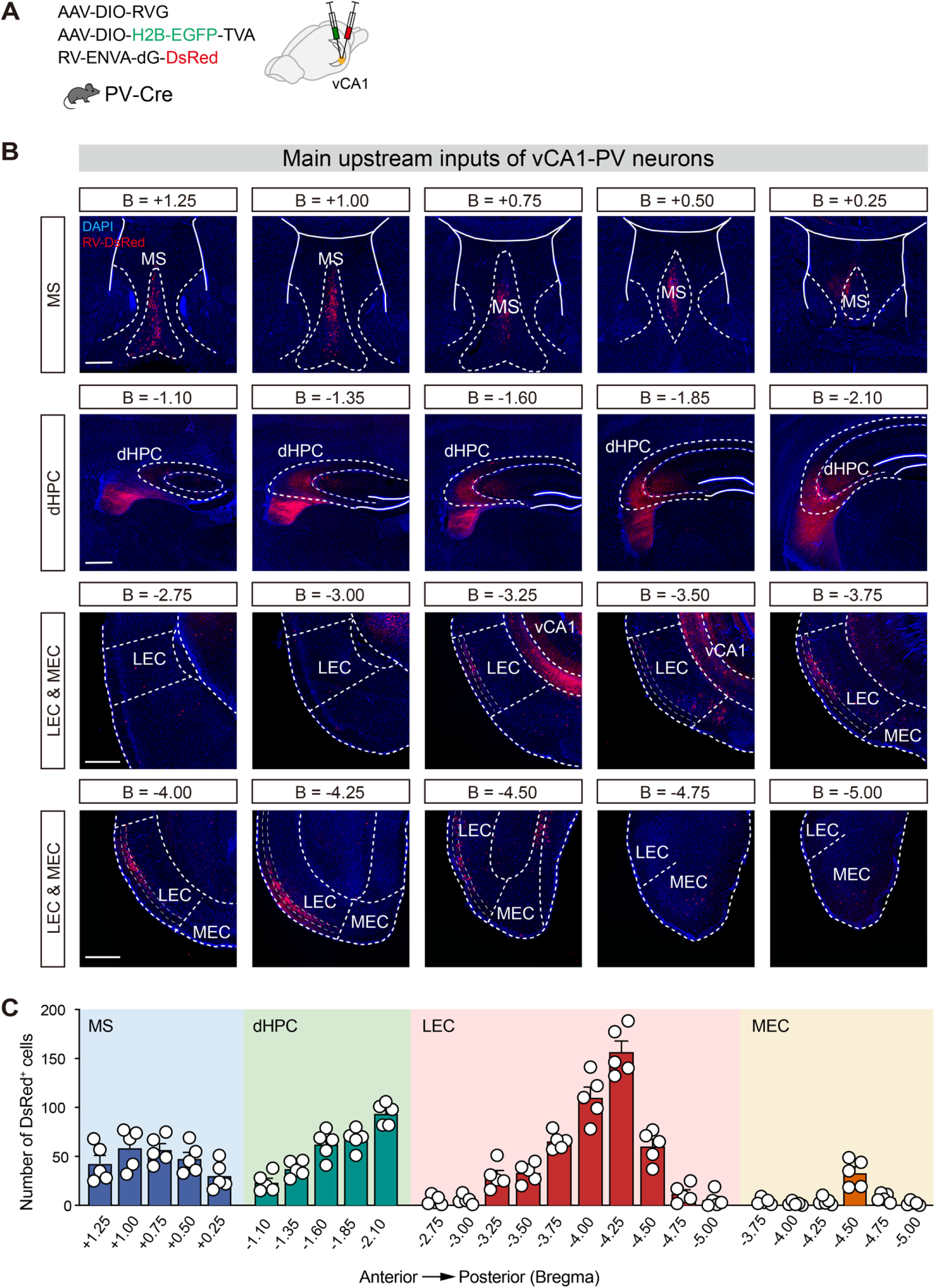
vCA1 PV projecting neurons at different bregma sites in MS, dHPC and EC. (**A**) Schematics of AAV injections and experimental design. (**B**) DsRed-labeled neurons at different bregma sites in MS, dHPC and EC of PV-Cre mice. Scale bar, 500 μm. (**C**) Data showing the number of DsRed-labeled cells at different bregma sites in MS, dHPC and EC of PV-Cre mice. Data are mean ± SEM. MS, medial septal nucleus; dHPC, dorsal hippocampus; LEC, lateral entorhinal cortex; MEC, medial entorhinal cortex.

**Figure S4.**
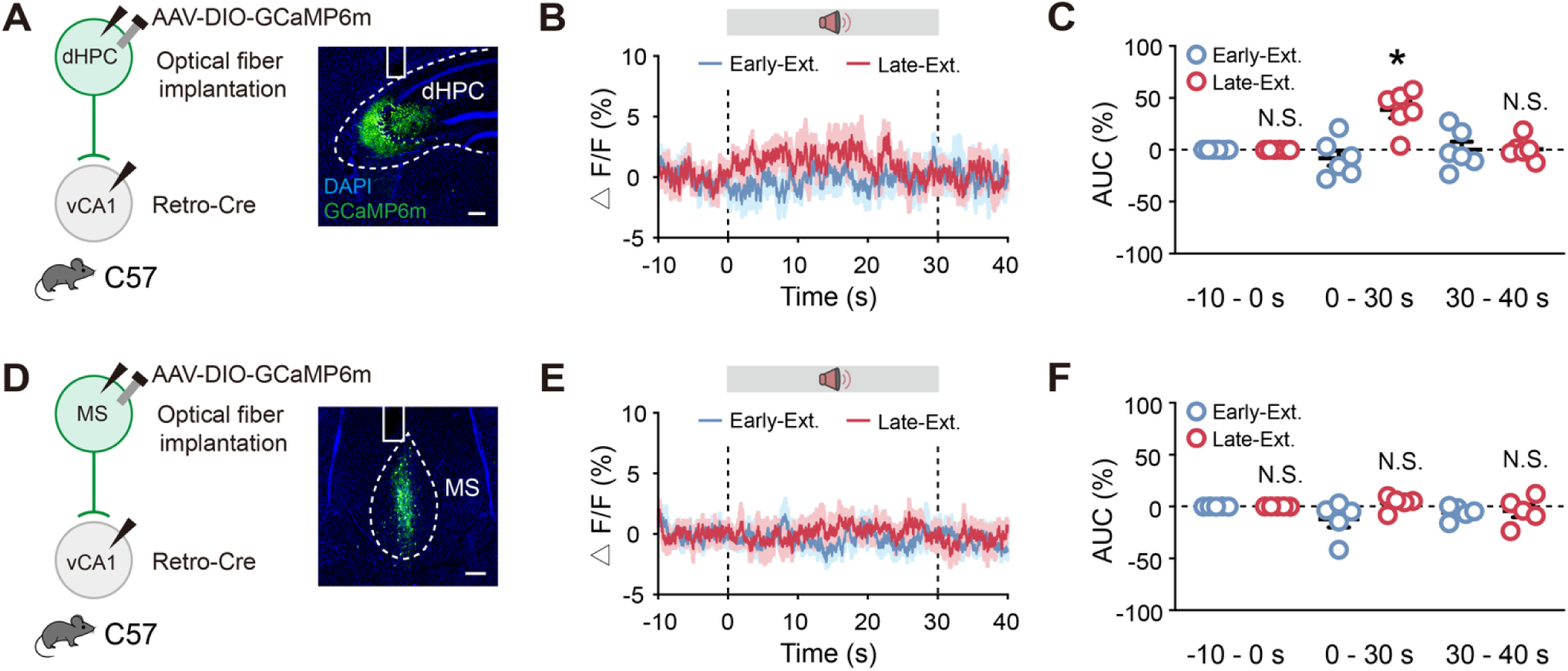
Ca^2+^ recordings of dHPC-vCA1 pathway and MS-vCA1 pathway during extinction. (**A**–**C**) Ca^2+^ recording of dHPC-vCA1 pathway during extinction. Schematics of AAV injections and fiber implantation (left). Representative images of CGaMP6m expression in dHPC (right). Scale bar, 200 μm (**A**). Average calcium signals during Early-Ext. and Late-Ext. (**B**). Activity of calcium signals (AUC) during Early-Ext. and Late-Ext. (**C**). Data are mean ± SEM. N.S., no significant difference, \**P* < 0.05, paired Student’s t-test. *n* = 5 mice. (**D**–**F**) Ca^2+^ recording of MS-vCA1 pathway during extinction. Schematics of AAV injections and fiber implantation (left). Representative images of CGaMP6m expression in MS (right). Scale bar, 200 μm (**D**). Average calcium signals during Early-Ext. and Late-Ext. (**E**). AUC during Early-Ext. and Late-Ext. (**F**). Data are mean ± SEM. N.S., no significant difference, paired Student’s t-test. *n* = 5 mice.

**Figure S5.**
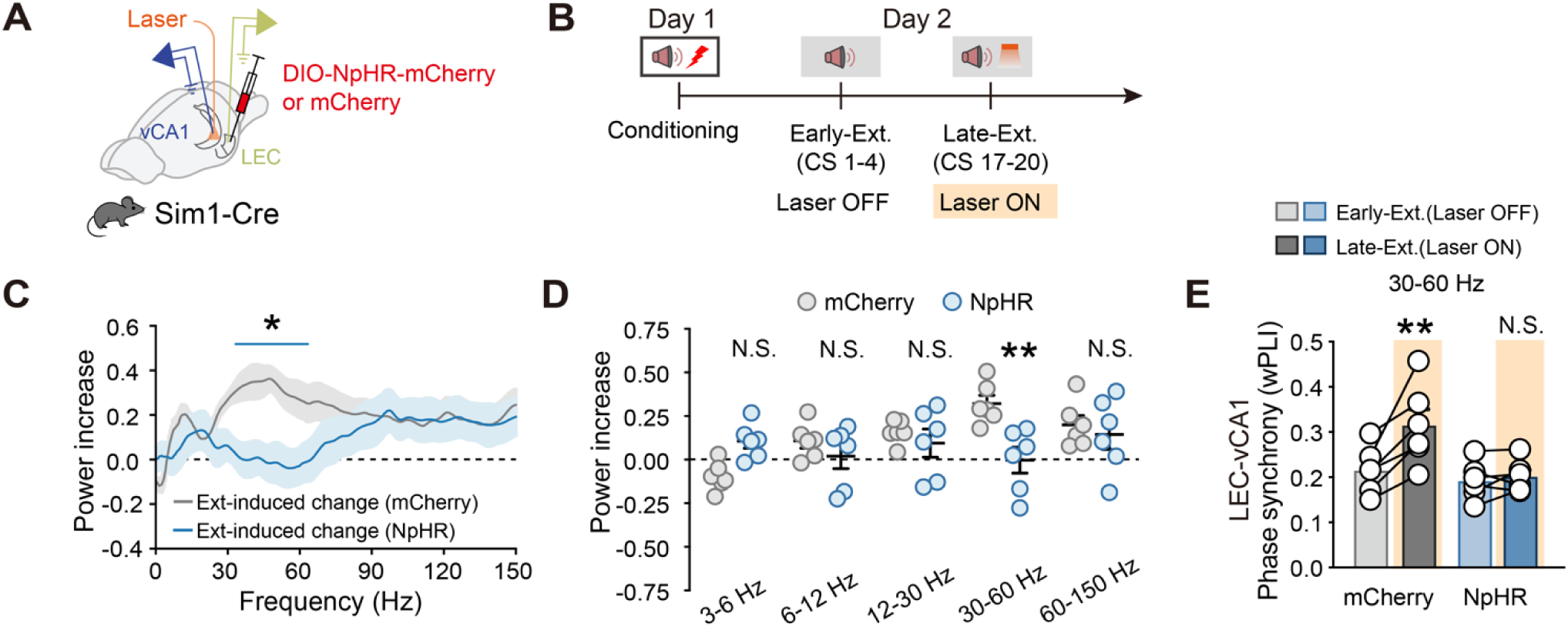
LEC-vCA1 projections promote network oscillations associated with fear extinction. (**A** and **B**) Schematics of stereotaxic surgery (**A**) and experimental design (**B**). Light for optogenetic inhibition was delivered during Late-Ext. on Day 2. (**C**) Extinction-induced changes for power spectrum of vCA1 LFP recordings. Mean ± SEM of power (Late-Ext. - Early-Ext.) / (Late-Ext. + Early-Ext.). *n* = 6 mice per group. Blue line indicates frequencies with a significant effect. \**P* < 0.05 with Bonferroni correction for multiple comparisons. (**D**) Average power increase of vCA1 LFP recordings. Data are mean ± SEM and circles denote individual mice. *n* = 6 mice per group. N.S., no significant difference, \*\**P* < 0.01, unpaired Student’s t-test. (**E**) Low-gamma phase synchrony quantified using the wPLI between LEC and vCA1 LFPs. Histograms represent mean ± SEM and circles denote individual mice. *n* = 6 mice per group. N.S., no significant difference, \*\**P* < 0.01, paired Student’s t-test.

**Figure S6.**
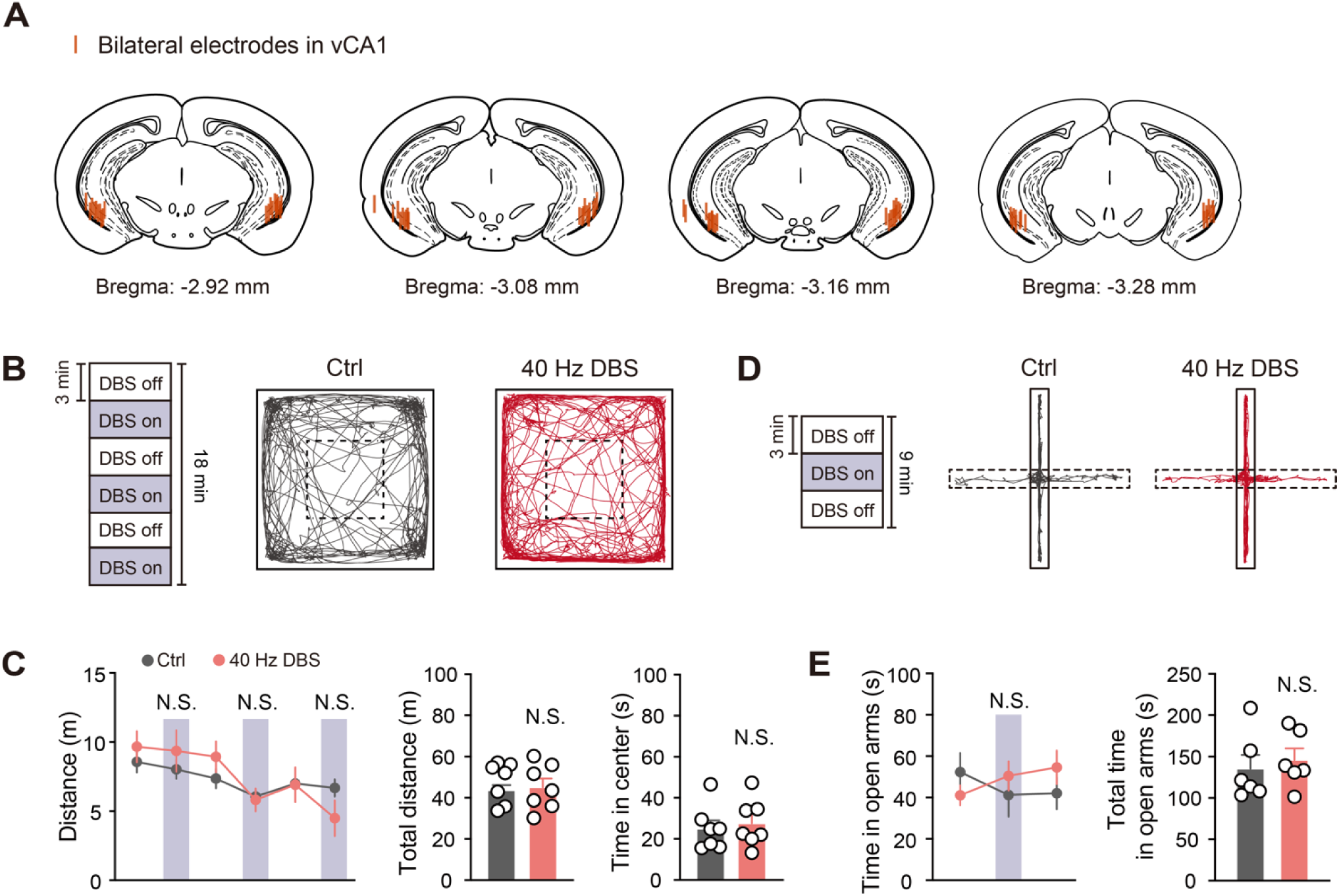
Low-gamma DBS-vCA1 had no effect on locomotor activity or anxiety-like behaviors. (**A**) Location of center for DBS electrode lesions for all mice. (**B** and **D**) Schematics of experimental design of the locomotor and anxiety-related activity for low-gamma DBS-vCA1. (**C**) Effect of low-gamma DBS-vCA1 on locomotor activity in the open field test (OFT). The distance traveled in the open field in a 3-min DBS-on (purple box) and DBS-off period (left). Total distances travelled (middle) and total time in center (right) in the open field for No DBS group and DBS group. Data are mean ± SEM. N.S., no significant difference, unpaired Student’s t-test. No DBS group, *n* = 7 mice; DBS group, *n* = 7 mice. (**E**) Effect of low-gamma DBS-vCA1 on anxiety-related activity in the elevated-plus maze (EPM). The time spent exploring open arms in a 3-min DBS-on (purple box) and DBS-off period (left). Total time spent exploring open arms (right) for No DBS group and DBS group. Data are mean ± SEM. N.S., no significant difference, unpaired Student’s t-test. No DBS group, *n* = 6 mice; DBS group, *n* = 6 mice.

**Figure S7.**
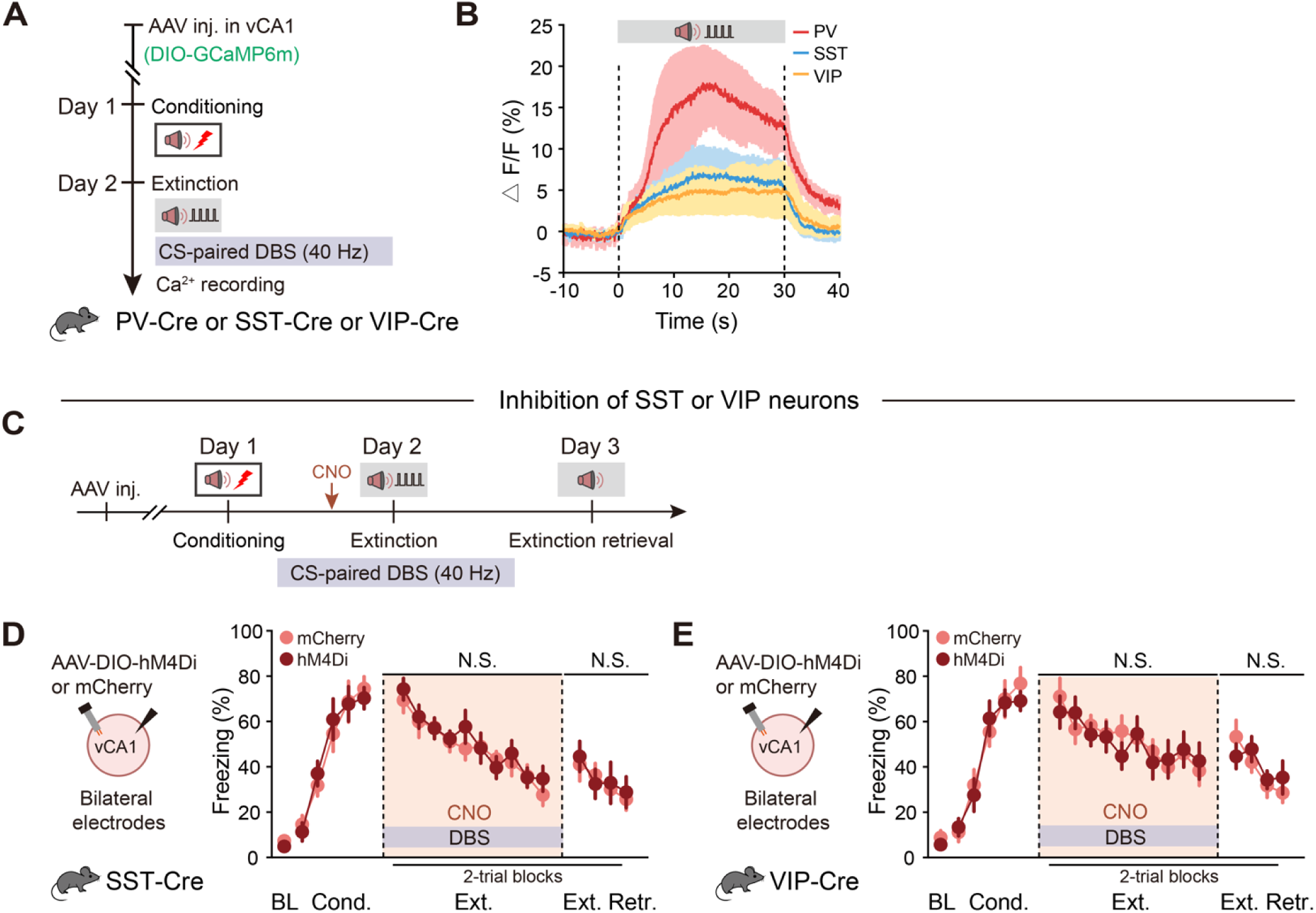
Low-gamma DBS enhanced PV-INs activity and inhibition of SST-INs or VIP-INs had no effect on DBS-induced extinction promotion. (**A** and **C**) Schematics of AAV injections and experimental design. (**B**) Average calcium signals in different INs during 40 Hz DBS-paired extinction training. Data are mean ± SEM. PV group, *n* = 4 mice; SST group, *n* = 4 mice; VIP group, *n* = 5 mice. (**D** and **E**) Effect of inhibiting vCA1 SST-INs (**D**) or VIP-INs (**E**) on DBS-induced extinction promotion. Schematics of AAV injections (left). Time courses of freezing responses to the CS during fear conditioning, extinction training and extinction retrieval sessions (right). Statistics are as follows: two-way repeated measures ANOVA, main effect of AAV, (**D**) conditioning, F_1,12_ = 0.0081, *P* = 0.9297; extinction training, F_1,12_ = 0.6232, *P* = 0.4452; extinction retrieval, F_1,12_ = 0.0374, *P* = 0.8499. mCherry group, *n* = 7 mice; hM4Di group, *n* = 7 mice. (**E**) conditioning, F_1,12_ = 0.0460, *P* = 0.8339; extinction training, F_1,12_ = 0.0661, *P* = 0.8015; extinction retrieval, F_1,12_ = 0.0788, *P* = 0.7837. mCherry group, *n* = 7 mice; hM4Di group, *n* = 7 mice. Data are mean ± SEM. N.S., no significant difference.

**Figure S8.**
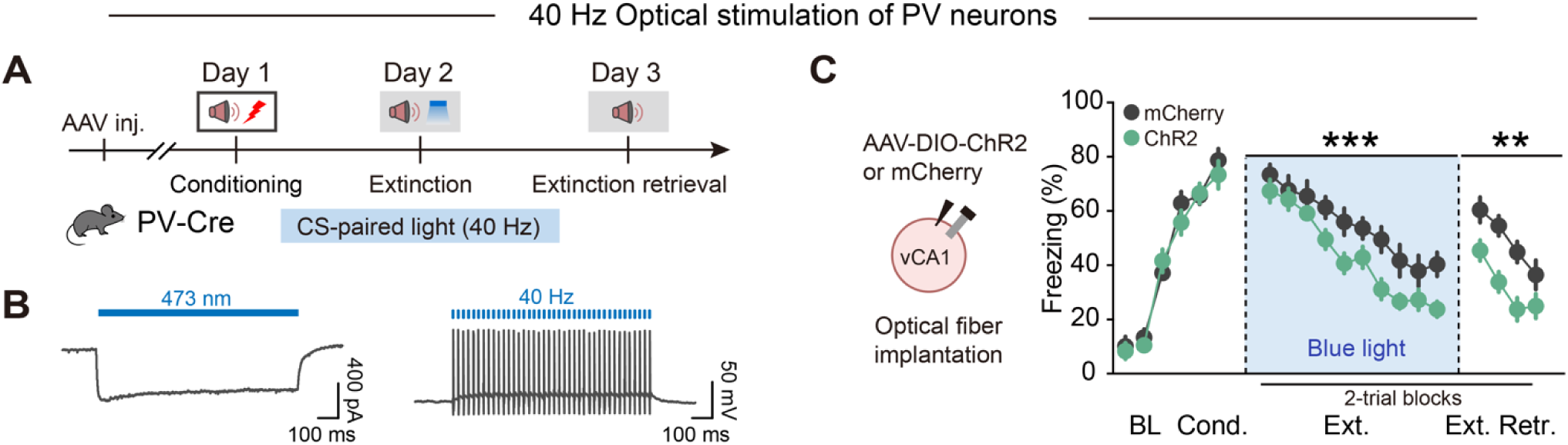
Effects of 40 Hz optical stimulation of PV neurons on extinction training and retrieval. (**A**) Experimental design of 40 Hz optical stimulation of vCA1 PV neurons. CS is paired with 40 Hz blue light during extinction training. (**B**) Activation effect of blue light on ChR2-expressing vCA1 PV neurons. Representative trace of ChR2-mediated current by 1-s pulses of blue light in the voltage-clamp mode (left). Action potentials were evoked in a ChR2-expressing vCA1 PV neuron in current-clamp mode by 1-ms pulses of photostimuli at 40 Hz (right). (**C**) Effects on optical activating vCA1 PV neurons on extinction training and extinction retrieval. Schematics of AAV injections (left). Time courses of freezing responses to the CS during fear conditioning, extinction training and extinction retrieval sessions (right). Statistics are as follows: two-way repeated measures ANOVA, main effect of AAV, conditioning, F_1,16_ = 0.6004, *P* = 0.4497; extinction training, F_1,16_ = 17.04, *P* = 0.0008; extinction retrieval, F_1,16_ = 15.35, *P* = 0.0012. mCherry group, *n* = 9 mice, ChR2 group, *n* = 9 mice. Data are mean ± SEM. N.S., no significant difference, \*\**P* < 0.01, \*\*\**P* < 0.001.

**Figure S9.**
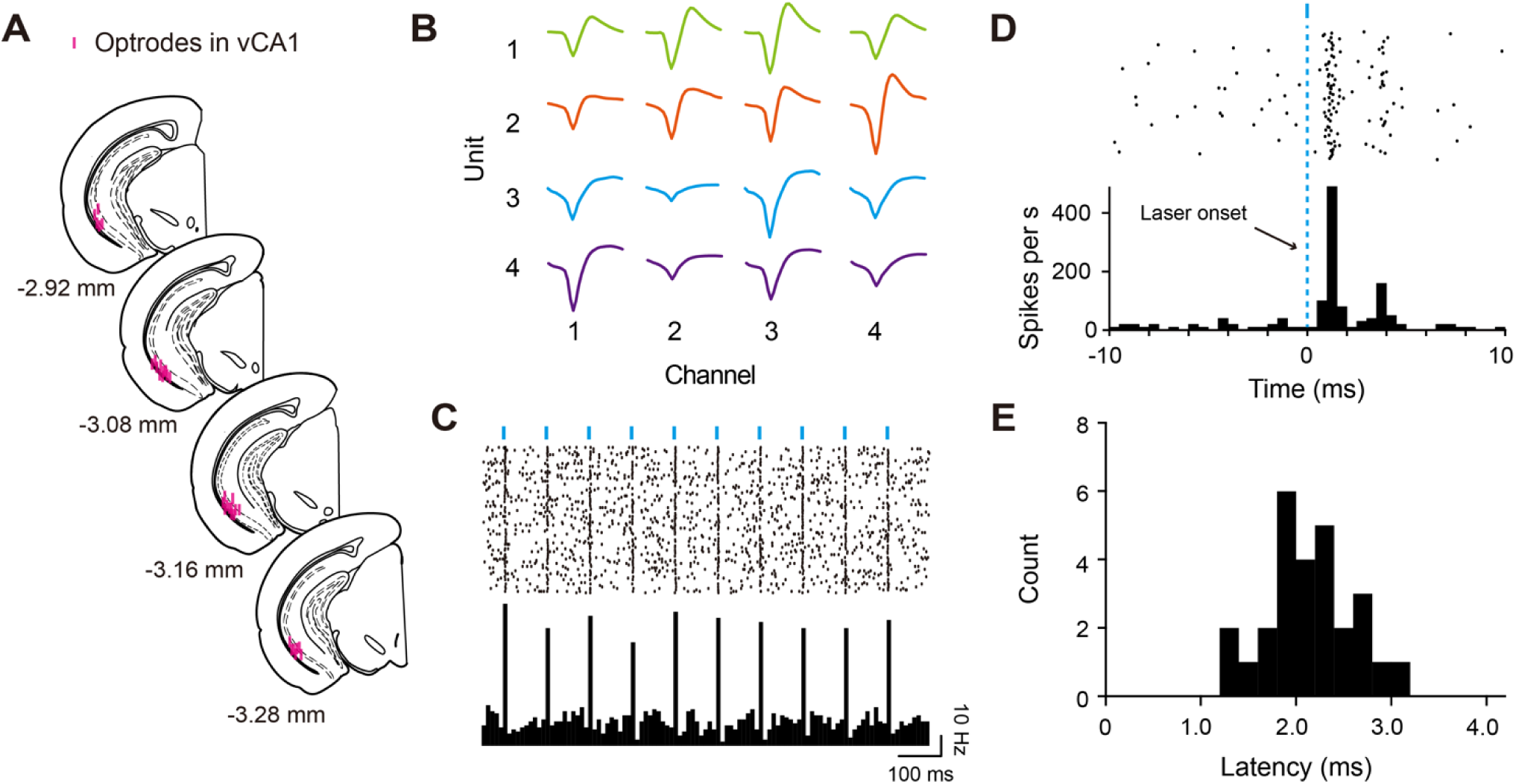
Single-unit recording of vCA1. (**A**) Optrode placement in vCA1 for optrode recording experiments. (**B**) An example spike sorting result from a single tetrode in vCA1. (**C**) Raster plot (top) and PSTH (bottom) for light-evoked spikes of an example tagged PV-IN. Blue bars mark light pulses (1-ms, 10 Hz). (**D**) Sample of peri-event raster plots and PSTH for light-evoked spike latency around 2 ms (left). Plot of latency distribution showing short latencies of light-evoked spike for all ChR2-tagged PV-INs in vCA1 (right). (**E**) Classification of recorded vCA1 neurons into WS putative pyramidal cells (blue circles), NS-nonFS (gray circles), Tagged PV (red circles) and FS-PV (orange circles) based on peak-to-trough latency and baseline firing rate.

**Figure S10.**
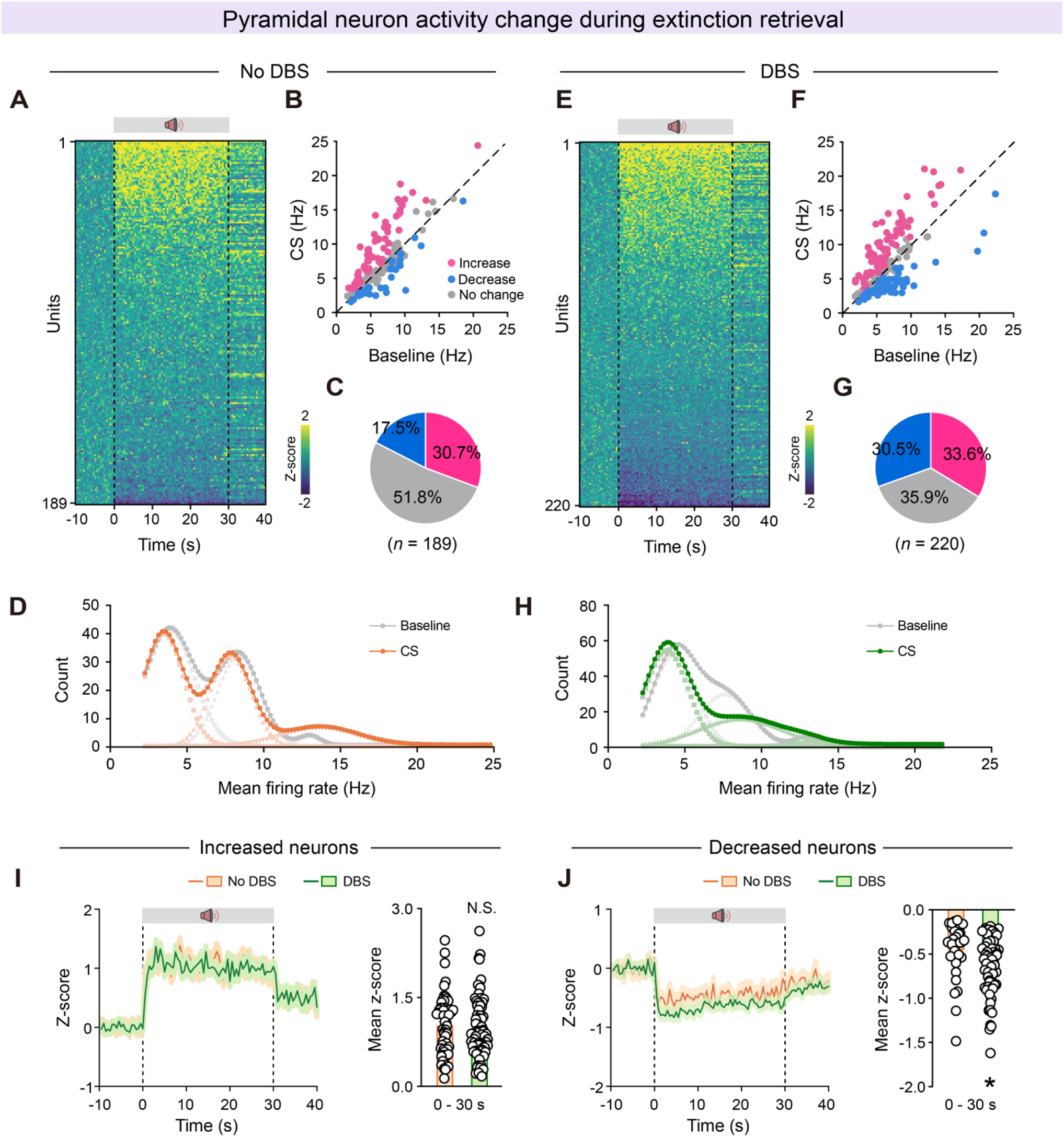
Low-gamma DBS paired extinction training induces sustained inhibition of vCA1 pyramidal neurons during extinction retrieval. (**A** and **E**) Heatmaps showing responses of pyramidal neurons during extinction retrieval. (**A**) No DBS manipulation during extinction training. (**E**) CS is paired with 40 Hz DBS during extinction training. (**B** and **F**) Correlation of firing rate during Baseline and CS for individual pyramidal neurons. (**B**) No DBS manipulation during extinction training. (**F**) CS is paired with 40 Hz DBS during extinction training. Colored circles indicate neurons that showed significant firing rate increase (pink) or decrease (blue) or no significant difference (gray). (**C** and **G**) The pie graphs show the percentage of neurons that had significantly higher, lower, or unchanged firing rates during extinction retrieval. (**D** and **H**) Distribution of mean firing rate during Baseline and CS for individual pyramidal neurons. (**D**) No DBS manipulation during extinction training. Gray lines indicate the distribution of mean firing rate during Baseline. Orange lines indicate the distribution of mean firing rate during CS. (**H**) CS is paired with 40 Hz DBS during extinction training. Gray lines indicate the distribution of mean firing rate during Baseline. Green lines indicate the distribution of mean firing rate during CS. (**I** and **J**) Z-scored signal changes of increased (**I**) and decreased (**J**) responses of pyramidal neurons during extinction retrieval. Orange lines indicate No DBS manipulation during extinction training and green lines indicate 40 Hz DBS manipulation during extinction training. Data are mean ± SEM. N.S., no significant difference, \**P* < 0.05, unpaired Student’s t-test.

**Figure S11.**
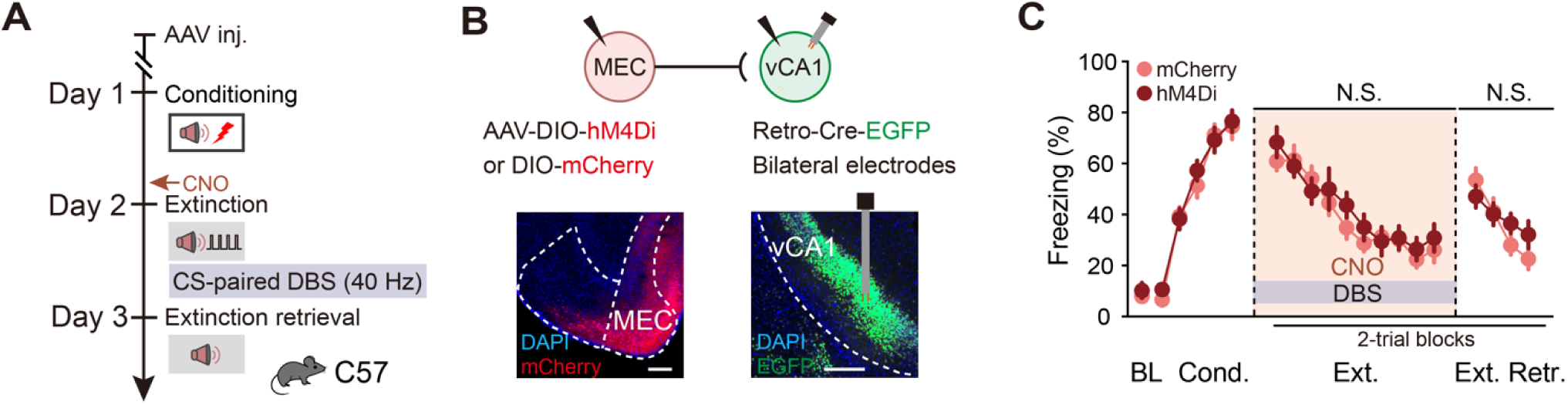
Inhibition of MEC-vCA1 had no effect on DBS-induced extinction promotion. (**A**) Schematics of experimental design. CS is paired with 40 Hz DBS during extinction training and CNO was administrated 30 min (i.p.) before extinction training. (**B**) Schematics of AAV injections (top) and representative images of virus expression (bottom). Scale bar, 200 μm. (**C**) Effect of inhibiting MEC-vCA1 projectors on DBS-induced extinction promotion. Time courses of freezing responses to the CS during fear conditioning, extinction training and extinction retrieval sessions. Statistics are as follows: two-way repeated measures ANOVA, main effect of AAV, conditioning, F_1,14_ = 0.4088, *P* = 0.5329; extinction training, F_1,14_ = 0.4952, *P* = 0.4932; extinction retrieval, F_1,14_ = 0.2920, *P* = 0.5974. mCherry group, *n* = 8 mice; hM4Di group, *n* = 8 mice. Data are mean ± SEM. N.S., no significant difference.

**Figure S12.**
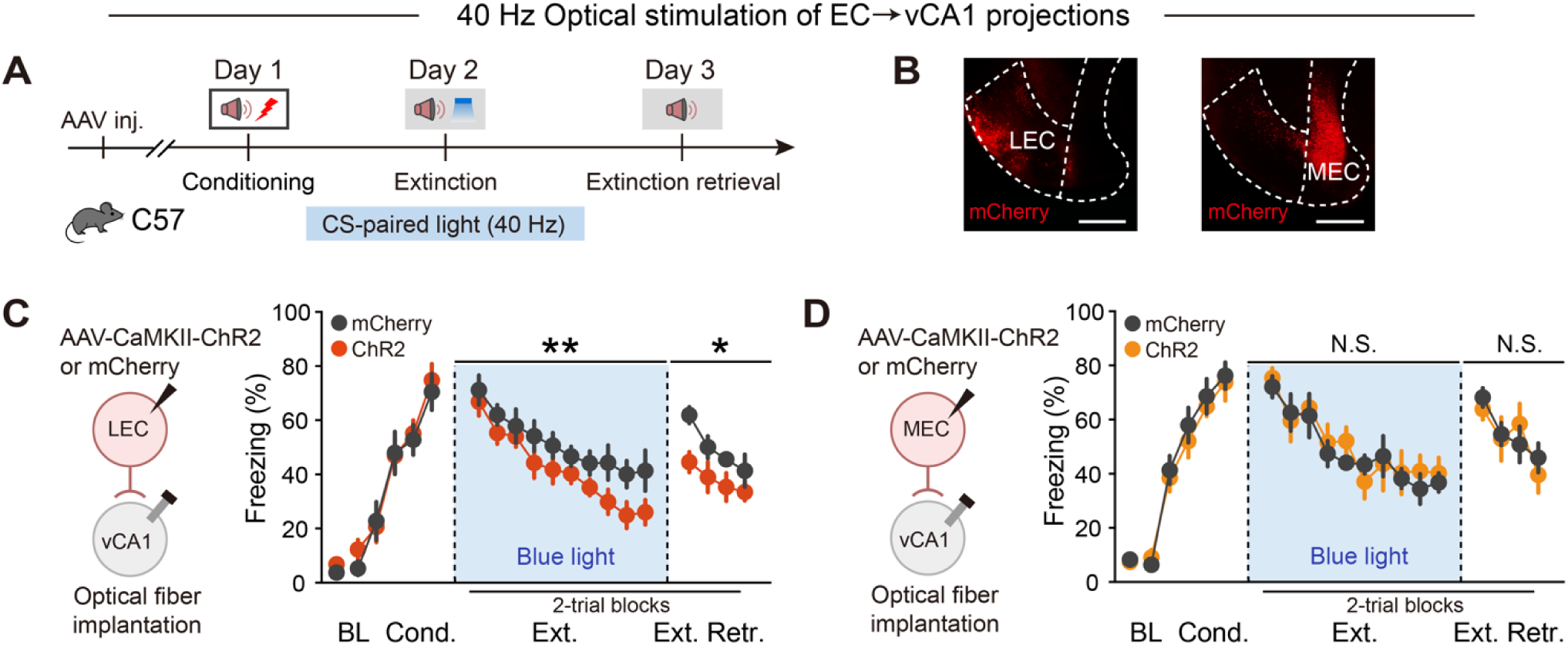
Effects of 40 Hz optical stimulation of LEC-vCA1 or MEC-vCA1 on extinction training and retrieval. (**A**) Experimental design of 40 Hz optical stimulation of EC→vCA1 projections. CS is paired with 40 Hz blue light during extinction training. (**B**) Representative images of virus expression. Scale bar, 500 μm. (**C** and **D**) Effects on optical activating LEC→vCA1 projection (**C**) and MEC→vCA1 projection (**D**) on extinction training and extinction retrieval. Schematics of AAV injections (left). Time courses of freezing responses to the CS during fear conditioning, extinction training and extinction retrieval sessions (right). Statistics are as follows: two-way repeated measures ANOVA, main effect of AAV, (**C**) conditioning, F_1,13_ = 0.1852, *P* = 0.6740; extinction training, F_1,13_ = 14.39, *P* = 0.0022; extinction retrieval, F_1,13_ = 7.201, *P* = 0.0188. mCherry group, *n* = 7 mice, ChR2 group, *n* = 8 mice. (**D**) conditioning, F1,12 = 0.2787, *P* = 0.6072; extinction training, F1,12 = 0.3980, *P* = 0.5399; extinction retrieval, F_1,12_ = 0.03806, *P* = 0.8486. mCherry group, *n* = 7 mice, ChR2 group, *n* = 7 mice. Data are mean ± SEM. N.S., no significant difference, \**P* < 0.05, \*\**P* < 0.01.

**Figure S13.**
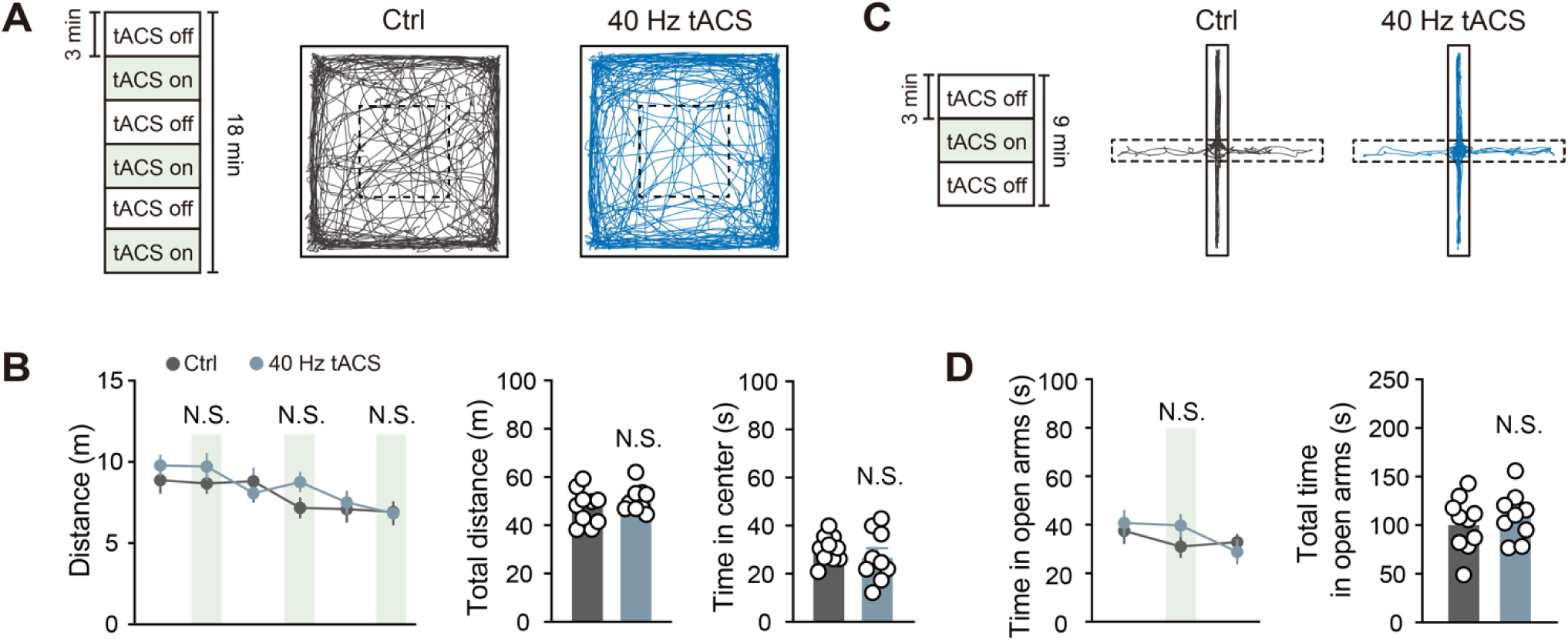
Low-gamma tACS-LEC had no effect on locomotor activity or anxiety-like behaviors. (**A** and **C**) Schematics of experimental design of the locomotor and anxiety-related activity for low-gamma tACS-LEC. (**B**) Effect of low-gamma tACS-LEC on locomotor activity in OFT. The distance traveled in the open field in a 3-min tACS-on (purple box) and tACS-off period (left). Total distances travelled (middle) and total time in center (right) in the open field for No tACS group and tACS group. Data are mean ± SEM. N.S., no significant difference, unpaired Student’s t-test. No tACS group, *n* = 10 mice; tACS group, *n* = 9 mice. (**D**) Effect of low-gamma tACS-LEC on anxiety-related activity in EPM. The time spent exploring open arms in a 3-min tACS-on (purple box) and tACS-off period (left). Total time spent exploring open arms (right) for No tACS group and tACS group. Data are mean ± SEM. N.S., no significant difference, unpaired Student’s t-test. No tACS group, *n* = 10 mice; tACS group, *n* = 9 mice.

## Notes

### Competing Interest Statement

The authors have declared no competing interest.

